# Bidirectional control of neurovascular coupling by cortical somatostatin interneurons

**DOI:** 10.1101/2024.11.06.622247

**Authors:** Boubacar Mohamed, Julie O’Reilly, Sandrine Picaud, Benjamin Le Gac, Esther Belzic, Liliana R V Castro, Hédi Soula, Isabelle Dusart, Dongdong Li, Bruno Cauli

## Abstract

Neurovascular coupling, linking neuronal activity to cerebral blood flow, is altered early in neurological disorders and underlies functional brain imaging. This process involves numerous cellular players. Among them inhibitory interneurons receive increasing attention, but how they control blood flow remains elusive. This study elucidates the mechanisms by which somatostatin interneurons bidirectionally control neurovascular coupling. Patch clamp recordings in, ex vivo, cortical slices from mice expressing channelrhodopsin-2 in somatostatin interneurons, revealed that these neurons are supralinearly activated at low-frequencies (< 5 Hz) and efficiently photostimulated at frequencies up to 20 Hz. Ex vivo vascular imaging showed that low-frequency (2 Hz) photostimulation triggered vasodilation whereas high-frequency (20 Hz) photostimulation induced vasoconstriction. Histochemistry revealed that subpopulations of cortical somatostatin interneurons expressed the neuronal nitric oxide synthase, and/or neuropeptide Y to a greater extent. Consistently, pharmacological investigations showed that vasodilation induced by low-frequency photostimulation involves nitric oxide release and activation of soluble guanylate cyclase. In contrast, the vasoconstriction induced at high-frequency photostimulation involves neuropeptide Y release and activation of the Y1 vascular receptor. These findings provide valuable insights into neurovascular coupling and help to understand the cellular mechanism underlying the functional brain imaging signals.

## INTRODUCTION

Brain function and integrity are critically dependent on a permanent blood supply provided by a dense vascular network (Schaeffer and Iadecola, 2021). Cerebral blood flow (CBF) is also tightly controlled by neuronal activity, an essential process called neurovascular coupling (NVC) that is impaired in neurological disorders (Iadecola, 2017). The NVC is thought to ensure the optimal supply of metabolic needs to, and removal of waste products and heat from the activated brain regions, although its precise motive remains to be understood (Drew, 2022). It allows CBF to surge within seconds of increased neuronal activity and serves as the physiological basis for functional brain imaging techniques to map brain function (Iadecola, 2017). In the cerebral cortex, this hyperemic response is supported by dynamically controlled vasodilation that is spatially and temporally confined by vasoconstriction (Devor et al., 2007; Boorman et al., 2010; Uhlirova et al., 2016).

CBF is critically orchestrated by pial and penetrating arterioles, which have a higher density of contractile mural cells and control their diameter more rapidly than capillaries (Hill et al., 2015; Rungta et al., 2018; Hartmann et al., 2021). NVC is mediated by multiple vasoactive agents produced by excitatory neurons, astrocytes, GABAergic interneurons, and/or endothelial cells (Cauli and Hamel, 2018; Kaplan et al., 2020; Schaeffer and Iadecola, 2021) that control vessel diameter and CBF along the vascular tree (Rungta et al., 2018). These agents may be vasodilatory [e.g. arachidonic acid derivatives, nitric oxide (NO)], vasoconstrictive [e.g. Neuropeptide Y (NPY) (Uhlirova et al., 2016)], or both [e.g. glutamate, K^+^ (Girouard et al., 2010; Zhang et al., 2024)].

The cellular diversity that characterizes cortical GABAergic interneurons (Ascoli et al., 2008), has made it challenging to study their contribution to NVC, and the conditions under which they release specific vasoactive agents have long remained elusive (Cauli et al., 2004). A major breakthrough in understanding their role came with the advent of transgenic mice and optogenetics, which allowed confirmation of their ability to induce vasodilation and/or vasoconstriction *in vivo* (Anenberg et al., 2015; Uhlirova et al., 2016). More recently the use of different Cre-line drivers has made it possible to investigate the contribution of different subpopulations of GABAergic interneurons (Krawchuk et al., 2019; Lee et al., 2020; Echagarruga et al., 2020; Krogsgaard et al., 2023; Ruff et al., 2024). Among these subclasses, somatostatin (Sst) interneurons have been found to either increase and/or decrease CBF depending on the stimulation paradigm (Krawchuk et al., 2019; Lee et al., 2020), but the underlying mechanisms of this bimodal response have not been determined.

To address this unresolved issue, we used *ex vivo* approaches in combination with optogenetics to precisely control Sst interneurons firing in the mouse barrel cortex while monitoring the resulting arteriolar response. We found that Sst interneurons are supralinearly activated by low-frequency (< 5 Hz) photostimulation and are efficiently and reliably activated up to 20 Hz. Low frequency stimulation (2 Hz) induced predominantly vasodilation, whereas 20 Hz photostimulation resulted in vasoconstriction. Histochemical analyses showed that subpopulations of Sst interneurons expressed the neuronal isoform of NO synthase, NOS-1, and/or NPY at much greater extents. Pharmacological investigations revealed that both types of vascular responses were dependent on action potential (AP) firing. The vasodilation required NO synthesis and activation of the soluble guanylate cyclase (sGC), whereas the vasoconstriction involved NPY and activation of its Y1 receptor. Thus, our study elucidates the mechanisms by which Sst interneurons can induce vasodilation or vasoconstriction depending on their firing frequency.

## MATERIALS AND METHODS

### Animal models

Homozygous Sst-IRES-Cre mice [Jackson Laboratory, B6J.Cg-*Sst^tm2.1(cre)Zjh^*/MwarJ, stock# 028864, (Taniguchi et al., 2011)] were crossed with homozygous Ai32 mice [Jackson Laboratory, B6.Cg-*Gt(ROSA)26Sor^tm32(CAG-COP4*H134R/EYFP)Hze^*/J stock #024109, (Madisen et al., 2012)] or with homozygous Ai95d mice [Jackson Laboratory, B6J.Cg-*Gt(ROSA)26Sor^tm95.1(CAG-GCaMP6f)Hze^*/MwarJ, stock# 028865 (Madisen et al., 2015)], to obtain heterozygous Sst^Cre/WT^::Ai32^ChR2/WT^ (Sst-ChR2) for optogenetic stimulations or Sst^Cre/Wt^::Ai95d^GCaMP6f/Wt^ (Sst-GCaMP6f) mice for immunohistochemistry. C57BL/6RJ mice were used for control optogenetic experiments and exogenous NPY applications. 16-21 days postnatal old females and males were used for all experiments. All experimental procedures involving animals were carried out in strict compliance with French regulations (Code Rural R214/87 to R214/130) and in accordance with the ethical guidelines of the Directive of the Council of the European Communities of September 22, 2010 (2010/63/EU).

### *Ex vivo* slice preparation

Mice were deeply anesthetized by isoflurane (IsoVet Piramal Healthcare UK or IsoFlo, Axience) evaporation in an induction box then euthanized by decapitation. The brain was quickly removed and immersed in cold (∼ 4°C) oxygenated artificial cerebrospinal fluid (aCSF) containing in (mM): 125 NaCl, 2.5 KCl, 1.25 NaH_2_PO_4_, 2 CaCl_2_, 1 MgCl_2_, 26 NaHCO_3_, 10 glucose, 15 sucrose and 1 kynurenic acid (Sigma-Aldrich). 300 µm-thick coronal slices containing the barrel cortex were cut with a vibratome (VT1000s; Leica) and were allowed to recover at room temperature for at least 45 min with oxygenated aCSF (95% O_2_/5% CO_2_) (Devienne et al., 2018). The slices were transferred to a recording chamber and continuously perfused at a rate of ∼2 ml/min with oxygenated aCSF lacking kynurenic acid.

### Patch-clamp recording

Borosilicate glass patch pipettes (5.0 ± 0.4 MΩ) were filled with an internal solution containing (in mM): 144 K-gluconate, 3 MgCl_2_, 0.5 EGTA, 10 HEPES, pH 7.2 (285/295 mOsm). Sst-ChR2 interneurons were first visually identified by their EYFP fluorescence. Whole-cell recordings were performed at room temperature (20-25 °C) using a patch clamp amplifier (Axopatch 200B, Molecular Devices) at a wavelength of 780 nm, which does not activate ChR2 (Lin et al., 2009). Data were filtered at 5-10 kHz and digitized at 50 kHz using an acquisition board (Digidata 1440, MDS) attached to a personal computer running pCLAMP 10.7 software package (MDS). Electrophysiological properties were determined using the fast I-clamp mode of the amplifier and were measured as previously described (Karagiannis et al., 2021, 2009). Membrane potential values were not corrected for liquid junction potential. Only neurons with a resting membrane potential more hyperpolarized than -55 mV were further analyzed.

### Optogenetic stimulation

Optogenetic stimulation performed through the objective using a 470 nm light-emitting diode (LED) device (CoolLED brand, Precise Excite model) attached to the epifluorescence port of a BX51WI microscope (Olympus) and a set of multiband filters consisting of an excitation filter (HC 392/474/554/635, Semrock), a dichroic mirror (BS 409/493/573/652, Semrock), and an emission filter (HC 432/515/595/730, Semrock). Photostimulation consisted of a 10-s train of 5 ms light pulses at an intensity of 38 mW/mm² and delivered at six different frequencies (1, 2, 5, 10, 20 and 40 Hz).

### Vascular imaging

Blood vessels and cells were observed in slices under infrared illumination with Dodt gradient contrast optics (IR-DGC Luigs and Neumann; (Dodt and Zieglgansberger, 1998)) using a collimated LED (780 nm; ThorLabs) as the transmitted light source, a 40X (LUMPlanF/IR, 40X/0.80 W, Olympus) and a digital camera (OrcaFlash 4.0LT+, Hamamatsu) attached to the microscope. Diving arterioles were selected by IR-DGC based on their well-defined luminal diameter (16.7 ± 0.5 µm, n = 144 arterioles), their length remaining in the focal plane of at least 50 µm, and the thickness of their wall (5.5 ± 0.1 µm, n = 144 arterioles). A resting period of at least 30 min (Lovick et al., 1999; Zonta et al., 2003) was observed after slice transfer. IR-DGC images were acquired at 0.1 Hz for pharmacological applications and at 1 Hz for optogenetic experiments using Imaging Workbench 6.1 software (Indec Biosystems). The focal plane was continuously maintained on-line using IR-DGC images of cells as anatomical landmark (Lacroix et al., 2015). After light-induced responses, arteriolar identity and reactivity were systematically verified by assessing their contractility (Hartmann et al., 2021) using the thromboxane A2 analog, U46619 (100 nM) (Cauli et al., 2004; Rancillac et al., 2006). Blood vessels that failed to constrict after this pharmacological application were discarded. Only one arteriole was monitored per slice.

### Double fluorescence immunolabeling

Mice were euthanized with pentobarbital (150 mg/kg, i.p., Euthasol Vet) and their brains were perfusion fixed (50 ml of ice-cold 4 % paraformaldehyde (PFA), in 0.1M phosphate buffer, pH 7.4), and post-fixed by immersion in 4 % PFA (2 h, 4°C). Brains were then cut in 50 μm-thick coronal sections with a VT-1000S vibratome (Leica). After washing in PBS, free-floating sections were blocked for 2 h at room temperature in a PBS 1X/0.25% Triton X-100/0,2% gelatin solution before being incubated overnight at 4°C with primary antibodies. Sections from 3 to 4 different mouse brains were simultaneously incubated overnight with the primary antibodies against GFP and specific markers. Antibodies included chicken anti-GFP (1:1000; Aves Labs GFP-1020, (Tricoire et al., 2010)), rabbit anti-NOS-1 (1:1000, Millipore AB5380, (Tricoire et al., 2010)), sheep anti-NPY (1:2000, Abcam ab6173, (Lee et al., 2010)). The respective immunoreactions were visualized with the following secondary antibodies: goat anti-chicken Alexafluor 488 (1:1000, Invitrogen A11039), donkey anti-rabbit Alexafluor 555 (1:1000, Invitrogen A31572) donkey anti-sheep Alexafluor 555 (1:1000, Invitrogen A21436). Nuclear counterstaining was performed with 100 ng/ml DAPI (4’,6-diamidino-2-phenylindole; Invitrogen) solution in PBS for 20 min. Sections were mounted with Fluoromount G (SouthernBiotech) on gelatin-coated slides for visualization. Images of immunostained material were acquired and processed in Airyscan 2 mode using a LSM 980 confocal microscope with a 20x objective (N Plan-Apochromat 20x) and ZEN Blue 3 software (Zeiss). Cell counting was achieved using the cell counter plug-in of FIJI software.

### Drugs

All pharmacological compounds were bath-applied after a 5-minute control period (aCSF) and vascular dynamics were recorded during bath application. The following drugs were dissolved in water: NPY (Enzo, ALX-163-003), Nω-Nitro-L-arginine methyl ester hydrochloride (L-NAME, Sigma-Aldrich, N5751), BIBP3226 (1 µM, Tocris, 2707). Tetrodotoxin (TTX, 1 µM, Latoxan, L8503) was dissolved in 90% acetic acid. 1H-[1,2,4] oxadiazolo[4,3-a]quinoxalin-1-one (ODQ, Sigma-Aldrich, O3636) was dissolved in DMSO and 9,11-dideoxy-9α,11α-methanoepoxy prostaglandin F2α (U-46619, 100 nM, Enzo, BML-PG023) in ethanol. Acetic acid, DMSO and ethanol doses were always used below 0.1% to avoid vascular interactions (Sun et al., 2010). NO pathway inhibitors, BIBP3226 and TTX were applied at least 30 min before optogenetic stimulation or exogenous NPY application.

### Vascular reactivity analysis

To compensate for potential x-y drifts, all images were realigned off-line using the “StackReg” plug-in (Thevenaz et al., 1998) of the ImageJ 1.54f software. Luminal diameter was measured on registered images using custom analysis software developed in MATLAB (MathWorks) (Lacroix et al., 2015). Only arterioles with a stable luminal diameter were analyzed further. Arterioles were considered stable if the relative standard deviation (RSD) of their luminal diameter during the baseline period was less than 5% (Lacroix et al., 2015). The impact of blockers, inhibitors, and antagonists on arterioles was evaluated by comparing the mean luminal diameter during the initial 5 minutes of the baseline period and the final 5 minutes of the treatment period prior to optogenetic stimulation or NPY application. Diameter changes (ΔD/D_0_) were expressed according to: (Dt – D_0_)/D_0_ where Dt is the luminal diameter at the time t and D_0_ the average diameter during the 5 minutes baseline period. Vasomotor responses and their parameters were determined from the diameter change traces [ΔD/D_0_ = f(t)] plotted using Clampfit 10.7 (MDS). To determine whether arterioles dilated or constricted, a Z-score was calculated from the diameter change traces using the formula: Z = (x - μ)/ σ, where both the mean μ and standard deviation σ were calculated using the values before photostimulation or NPY application. Vascular responses were classified as vasodilation or vasoconstriction, respectively, if the Z-score exceeded a value of 1.96 (95% criteria) or fell below -1.96 for 10 seconds and 1 minute for photostimulation and NPY application, respectively. For each vascular response, the time of onset and peak (relative to the beginning of stimulation), duration, maximal amplitude and magnitude (area under the curve [AUC]) were determined (Supplementary Fig. 2). When multiple events of vasodilation or vasoconstriction occurred, the earliest onset was used for global onset, the time and amplitude of the largest maximum amplitude were used for global time to peak and global maximum amplitude, and the sum of durations and magnitudes was used for global duration and magnitude.

### Statistical analyses

Statistical analyses were performed using GraphPad Prism version 8.0.2 (GraphPad Software, La Jolla, California, USA). All values are expressed as means ± SEM. Normality of distribution was assessed using the Shapiro-Wilk test. Parametric tests were only used if this criterion was met. The mean spike success rate per light pulse between photostimulation frequencies was compared using the Q Friedman test. Comparisons of vascular responses were determined using Brown-Forsythe ANOVA or Kruskal-Wallis tests. Given p-values were corrected for multiple comparisons using the original False Discovery Rate method of Benjamini and Hochberg (Benjamini and Hochberg, 1995). Statistical significance on all figures uses the following convention: *p < 0.05, **p < 0.01 and ***p < 0.001.

## RESULTS

### Sst interneurons are efficiently activated by optogenetic stimulation over a wide range of frequencies

To evaluate the frequency dependence of the vascular response driven by Sst interneurons, we optogenetically elicited their AP firing while visualizing the evoked vascular response in cortical brain slices from Sst^Cre/WT^::Ai32^ChR2/WT^ mice (Madisen et al., 2012; Taniguchi et al., 2011) conditionally expressing the H134R variant of channelrhodopsin-2 (ChR2) in Sst interneurons. We first determined the electrophysiological phenotype of Sst-ChR2 cortical neurons using whole-cell patch clamp recordings. Cortical neurons expressing EYFP/ChR2 were selected based on their EYFP fluorescence (Fig. 1a and Supplementary Fig. 1) in layers II-III (n=13 neurons) and V (n=3 neurons). Most neurons exhibited electrophysiological features typical of adapting-Sst interneurons (Karagiannis et al., 2009, 2021; Perrenoud et al., 2012a), including a relatively high membrane resistance (608 ± 231 MΛ) and a pronounced voltage sag (21.2 ± 10.8 %) in response to hyperpolarizing current pulses (Fig. 1a, middle traces). They also exhibited a low rheobasic current (8 ± 14 pA, Fig. 1a) and a slowly developing early frequency adaptation (adaptation time constant: 60.5 ± 23.8 ms) when depolarized by high stimulation intensities (Fig. 1a). Consistently, the hyperpolarization (AHP) phase of their action potentials consisted of a fast AHP, followed by an afterdepolarization (ADP) and a medium AHP (Fig. 1a and Supplementary Table 1). However, as previously reported in Sst-Cre cortical neurons (Hu et al., 2013) and expected from the occasional expression of Sst transcripts in parvalbumin neurons (Cauli et al., 1997; Tasic et al., 2016), one Sst-ChR2 neuron exhibited the electrophysiological features of a fast spiking cell with a low input resistance (129 MΛ), a fast time constant (11.7 ms), a high rheobase (185 pA), short spikes (0.6 ms) with a sharp fast AHP (-26.5 mV), and the ability to fire high firing rates (maximal frequency: 99 Hz) with little frequency adaptation (Supplementary Fig. 1 and Table 1).

**Figure 1:**
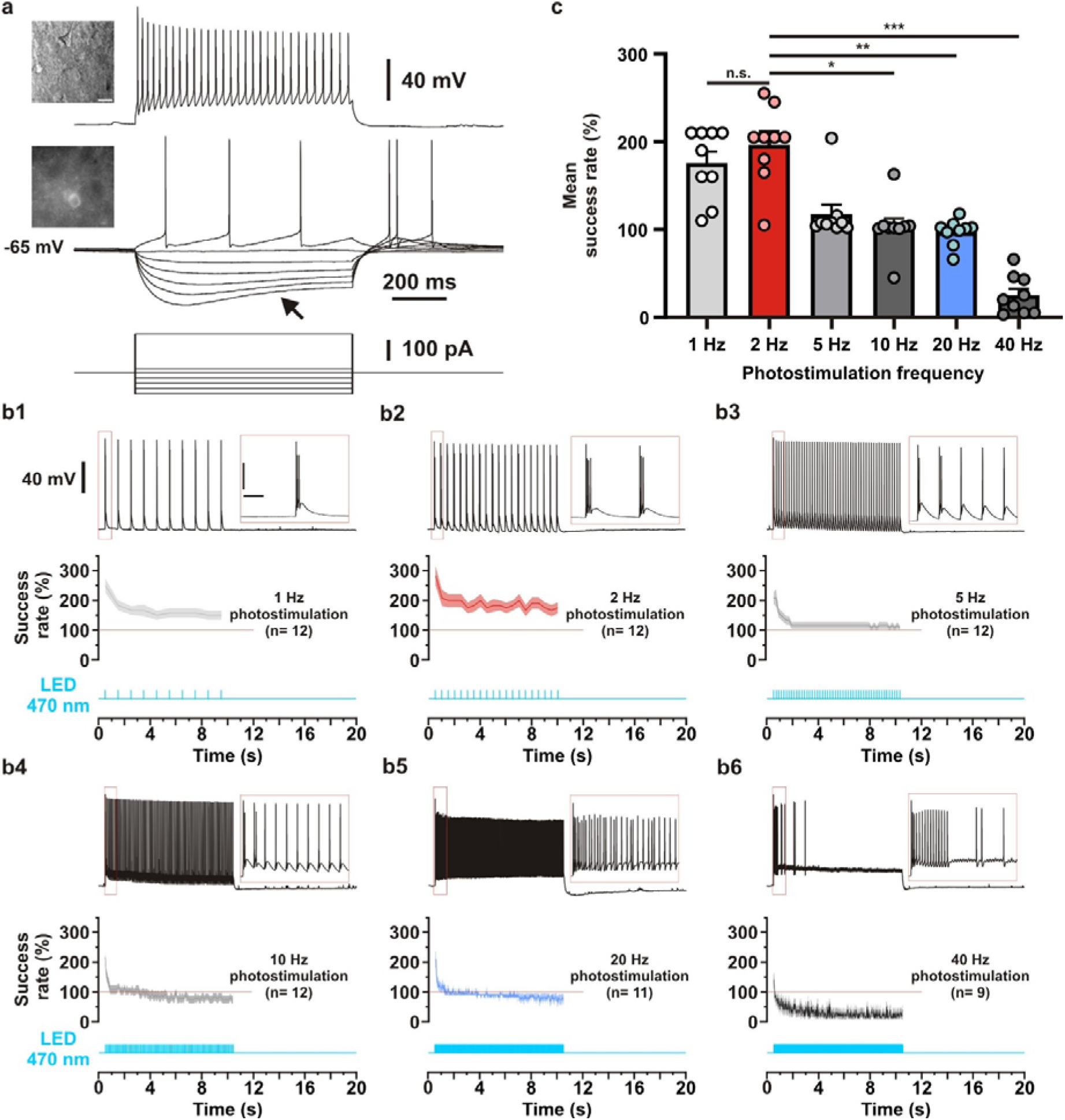
Electrophysiological characterization of cortical Sst-ChR2 neurons. **(a)** Representative recording showing the voltage responses induced by 800 ms current injections (bottom traces) of - 100 pA, -80 pA, -60 pA, -40 pA, -20 pA, 0 pA, +15 and +150 pA in a cortical EYFP/ChR2-expressing neuron from a Sst-Cre:Ai32 mouse. Note the pronounced voltage sag after the initial peak response (arrow) to hyperpolarizing current injections (middle traces). Just-above threshold current pulse (+15 pA) triggered a discharge of three action potentials (middle traces). Note the complex AHP consisting of a fast AHP, a small ADP, and a medium AHP. Near saturation, a strong depolarizing current (+150 pA) elicited a discharge of action potentials with a marked frequency adaptation and a monotonous amplitude accommodation (upper trace). Top inset: IR videomicroscopy picture of the recorded neuron, the pial surface is upward (scale bar, 20 µm). Bottom insets, corresponding field of view showing the EYFP fluorescence of the recorded neuron. **(b1-6)** Voltage responses (upper traces) of the Sst-ChR2 neuron shown in **(a)** induced by photostimulations (470 nm, 10 s train, 5 ms pulses) delivered at 1, 2, 5, 10, 20 and 40 Hz (cyan lower traces) and instantaneous spike success rate (middle trace). The red boxes highlight the first second of the zoomed recording in the right insets (scaled bars 200 ms, 40 mV). The red lines represent a spike success rate of 100%. Note that more than one action potential is elicited per light pulse delivered at 1 or 2 Hz whereas at frequencies above 5 Hz this success rate rapidly decreases to a maximum of one spike per light pulse. (**c**) Histogram showing the average spike success rate induced by photostimulations at 1, 2, 5, 10, 20, or 40 Hz (n = 9 Sst-ChR2 cortical neurons from 6 SST-Cre:Ai32 mice). Note the average spike success rate exceeding 100 % at photostimulation frequencies lower than 10 Hz. Data are presented as means ± SEM. The SEMs envelope the mean traces Statistically different with *p< 0.05, **p< 0.01 and ***p< 0.001; n.s.: not statistically significant.

Next, we evaluated the efficiency of optogenetic stimulation by recording the voltage response of the patched Sst-ChR2 neurons. Wide-field photostimulation was achieved using 10-second trains of 5 ms light pulses (see Materials and Methods) delivered at six different frequencies (1, 2, 5, 10, 20 and 40 Hz, Fig. 1b, lower traces). Surprisingly, at the beginning of the light pulse trains, Sst-ChR2 neurons fired up to five APs per light pulse (Fig. 1b, upper traces). Such bursting response declined within seconds before reaching a steady state level (Fig. 1b, middle traces). The instantaneous spike success rate remained above 100 % at stimulation frequencies below 10 Hz, while it decreased below 100% at higher frequencies (Fig. 1b, middle traces). Consistent with the kinetic properties of the H134R ChR2 variant (Lin et al., 2009), Sst-ChR2 neurons were not able to sustain the firing of action potentials when photostimulated at 40 Hz (Fig. 1b6). The mean spike success rate over the 10 s train differed between photostimulation frequencies (Fig. 1c, Q_(6,9)_= 41.89, p= 6.203 10^-8^, 1 Hz: 176 ± 13%; 2 Hz: 197 ± 15%; 5 Hz: 118 ± 11%; 10 Hz: 103 ± 10%; 20 Hz: 97 ± 5%; 40 Hz: 25 ± 7%, n=9 Sst-ChR2 neurons). Indeed, photostimulation at 1 Hz and 2 Hz showed a higher number of APs per light pulse than that observed at frequencies above 5 Hz (Fig. 1c, 1Hz vs. 10 Hz, p= 0.0374; 1Hz vs. 20 Hz, p= 0.0063; 1 Hz vs. 40 Hz, p= 7.000 10^-6^; 2 Hz vs. 10 Hz, p = 0.0245; 2Hz vs. 20 Hz, p= 0.0039, 2 Hz vs. 40 Hz, p=3.597 10^-6^). Taken together, these observations indicate that optogenetic stimulation efficiently and reliably activates Sst-ChR2 neurons, mostly of the adapting Sst subtype, at frequencies up to 20 Hz. They also reveal that these neurons over-respond to low frequencies of photostimulation.

### The polarity of the vascular response induced by the optogenetic stimulation of Sst-ChR2 neurons is frequency dependent

Next, we visualized the optogenetically induced response of diving arterioles in cortical slices (Fig. 2). We focused on the photostimulations at 2 Hz and 20 Hz, which correspond respectively to the highest AP number per light pulse and to the frequency that produced the highest total AP number, respectively (Fig. 1b,c). When photostimulation was delivered at 2 Hz in Sst-ChR2 slices, we observed that the vast majority of arterioles exhibited vasodilatory events (n= 13 of 14 arterioles, Fig. 2a,b), which could be accompanied by vasoconstrictive events (n=6 of 14). On average, this resulted in a slight vasodilation (Fig. 2c). High light intensity can cause dilation (Rungta et al., 2017) or constriction (Choi et al., 2010) independent of ChR2, so we did control experiments by repeating 2 Hz photostimulation in slices of C57Bl6 mice that do not express ChR2 (Fig. 2c). Although the overall vascular response, as estimated by the area under the curve (AUC) of the response following the optogenetic stimulation did not differ between Sst-ChR2 (55.8 ± 24.1 %.min, n= 14 arterioles) and C57Bl6 arterioles (37.4 ± 15.3 %.min, n= 6 arterioles, p= 0.15146, Fig. 2g1), an evident tendency was observed toward vasodilation in Sst-ChR2 arterioles (Fig. 2c). Because both vasodilation and vasoconstriction can occur in the same arteriole (Fig. 2b), we refined the analysis of vascular responses by separating vasodilatory events from vasoconstrictive events based on a Z-score test (supplementary Fig. 2a-b, see Methods). The magnitude of vasodilation was greater in Sst-ChR2 arterioles (69.6 ± 19.9 %.min, Fig. 2g2) than in control arterioles (8.5 ± 4.5 %.min, p=0.03415), although not the maximal amplitude (Sst-ChR2: 6.1 ± 1.3 % *vs.* C57Bl6: 2.2 ± 0.7 %, p= 0.12763, Fig. 2g3). Vasodilation also occurred earlier and lasted longer in Sst-ChR2 arterioles (onset: 221 ± 130 s, duration: 1121 ± 195 s) than in control arterioles (onset: 1013 ± 314 s, p= 0.02501, Fig. 2g4, duration: 304 ± 135 s, p= 0.04865, Fig. 2g6) but did not differ in their time to peak (Sst-ChR2: 669 ± 155 s *vs.* C57Bl6: 1014 ± 314 s, p= 0.32174, Fig. 2g5). In contrast, the amplitude, magnitude, and kinetic properties of vasoconstriction induced by photostimulation at 2 Hz did not differ between Sst-ChR2 and C57Bl6 arterioles (supplementary Fig. 2c1-5).

**Figure 2:**
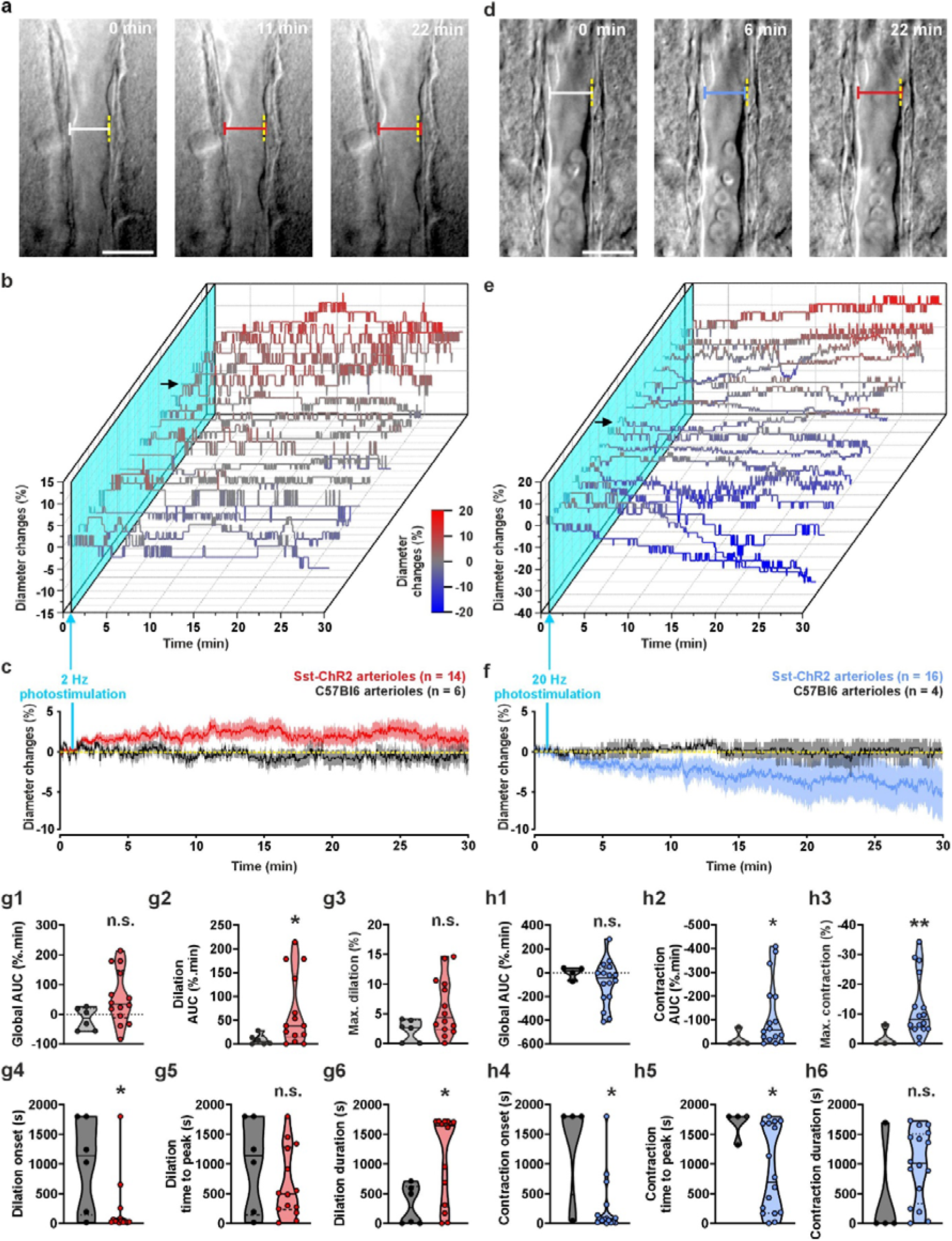
Bidirectional frequency-dependent vascular response induced by optogenetic stimulation of cortical Sst-ChR2 neurons. **(a, d)** Representative examples of infrared pictures of diving arterioles and diameter measurements before (0 min) and after (6, 11, 22 min) photostimulation at 2 Hz **(a)** and 20 Hz **(d)** in Sst-Cre:Ai32 cortical slices. The pial surface is upward. Scale bars 20 µm. **(a, d)** White, red and blue calipers correspond to baseline, dilated, and contracted arteriolar diameters, respectively. The dashed yellow lines indicate the initial diameters. Note the slight increase **(a)** and decrease followed by an increase **(d)** in diameter induced by photostimulation delivered at 2 Hz **(a)** and 20 Hz **(d)**, respectively. **(b, e)** Waterfall plots showing the kinetics of changes in diameter of single arterioles induced by photostimulation at 2 Hz **(b)** and 20 Hz **(e)**. The inset shows the color code used for constriction (blue) and dilation (red) used in both panels. Arterioles are sorted by increasing values of total vascular changes. The cyan planes and arrows indicate the onset of photostimulation. Black arrows indicate changes in diameter of the arterioles shown in **(a)** and **(b)**. **(c, f)** Kinetics of mean diameter changes induced by 2 Hz **(c)** or 20 Hz **(f)** photostimulation. The dashed yellow lines correspond to the initial diameters. **(c)** Note the average small diameter increase induced by 2 Hz photostimulation in Sst-Cre:Ai32 (red, n= 14 arterioles from 8 mice) but not in C57Bl6 arterioles (black, n= 6 arterioles from 5 mice) and **(f)** the average progressive diameter decrease induced by 20 Hz photostimulation in Sst-Cre:Ai32 (blue, n= 16 arterioles from 10 mice) but not in WT arterioles (black, n= 4 arterioles from 3 mice). **(g1-6, h1-6)** Violin plots summarizing the comparison of overall vascular response **(g1, h1)**, magnitude of dilation **(g2)** and contraction **(h2)**, maximum dilation **(g3)** and contraction **(h3)**, onset **(g4,h4)** time to peak **(g5,h5)** and duration **(g6,h6)** of dilation **(g2-6)** and contraction **(h2-6)** between arterioles from slices of Sst-Cre:Ai32 and C57Bl6 mice stimulated at 2 **(g1-6**) and 20 Hz **(h1-6)**. Note the larger **(g2),** earlier **(g4)** and longer **(g6)** dilation in Sst-ChR2 slices stimulated at 2 Hz and the larger **(h2,3)**, and earlier onset **(h4)** and time to peak **(h5)** contraction in Sst-ChR2 slices stimulated at 20 Hz Data are presented as individual values. Solid and dashed black bars correspond to median and quartile values. * and ** statistically different from C57Bl6 mice with p< 0.05 and 0.01, respectively. n.s.: not statistically significant.

We next evaluated whether a stronger optogenetic stimulation could induce a similar vascular response. Surprisingly, photostimulation at 20 Hz of Sst-ChR2 slices (n= 16) induced vasoconstrictive events in the vast majority of arterioles (n= 15 of 16), but also vasodilatory events to a lesser extent (n=9 of 16, Fig. 2d,e). This resulted on average in a progressive vasoconstriction that was not observed in control arterioles (n= 4, Fig. 2f). Although the overall vascular response was not statistically different between Sst-ChR2 and C57Bl6 arterioles (p=0.13458, Fig. 2h1), similarly to 2 Hz photostimulation it was more heterogeneous in Sst-ChR2 arterioles (-83.3 ± 46.7 %.min) than in C57Bl6 arterioles (0.0 ± 25.3 %.min) and may be associated with both vasoconstriction and vasodilation (Fig. 2d,e and supplemental Fig. 2 a,b). Both the magnitude of vasoconstriction and the maximal vasoconstriction were greater in Sst-ChR2 arterioles (AUC: -121.7 ± 35.4 %.min, Fig. 2h2; Maximal constriction: -12.2 ± 2.7 %, Fig. 2h3) than in control C57Bl6 (AUC: -17.8 ± 17.8 %.min, p=0.01810; Maximal constriction: -1.6 ± 1.6 %, p=0.00505). The onset and time to peak of vasoconstriction were also earlier in Sst-ChR2 arterioles (onset: 267 ± 120 s, Fig. 2h4; time to peak: 874 ± 181 s, Fig. 2h5) compared to control C57Bl6 arterioles (onset: 1361 ± 439 s; p= 0.01993; time to peak: 1680 ± 121 s, p= 0.04397). The contraction duration did not differ between Sst-ChR2 arterioles (981 ± 152 s) and C57Bl6 (424 ± 424 s, p= 0.13849, Fig. 2h6). In contrast, the amplitude, magnitude, and kinetic properties of vasodilatory events induced by photostimulation at 20 Hz did not differ between Sst-ChR2 and C57Bl6 arterioles (supplementary Fig. 2d1-5). Taken together, these observations indicate that, depending on the frequency of optogenetic stimulation Sst-ChR2 neurons elicit a bidirectional vascular response consisting mainly of vasodilation and vasoconstriction at 2 Hz and at 20 Hz, respectively.

### NOS-1 and NPY are expressed by subpopulations of Sst cortical interneurons

Subpopulation of Sst-expressing cortical interneurons have been reported to produce NO and/or NPY (Hendry et al., 1984; Kubota et al., 1994, 2011; Perrenoud et al., 2012a). These mediators may explain the bidirectional vascular responses induced by the photostimulation of Sst-ChR2 interneurons at 2 and 20 Hz, respectively, as they have been shown to induce vasodilation and vasoconstriction, respectively (Cauli et al., 2004; Rancillac et al., 2006). We therefore used double immunohistofluorescence to determine the extent to which Sst interneurons have the ability to produce NO and NPY. Consistent with previous reports (Kubota et al., 1994, 2011; Perrenoud et al., 2012a), NOS-1 immunolabeling revealed that strongly immunoreactive somata appeared denser in deep than in superficial cortical layers (Fig. 3a2-3 and supplementary Fig. 3). Some of these neurons were clearly identified as Sst-ChR2 interneurons as shown by immunolabelling of the ChR2-EYFP fusion transgene using an anti-GFP antibody (supplementary Fig. 3). However, consistent with the membrane localization of ChR2-EYFP (Fig. 1a and supplementary Fig. 1), GFP immunofluorescence was present at the periphery of the somata and in a dense fiber plexus (supplementary Fig. 3) making it difficult to evaluate the degree of co-expression. Therefore, we thus used Sst-GCamp6 mice (n= 3 mice) to obtain somatic Cre-dependent expression of the circularly permuted GFP of GCamp6 (Madisen et al., 2015). As previously reported in Sst-IRES-Cre mice (Taniguchi et al., 2011), Cre-dependent GFP expression was observed in the somata of neurons distributed throughout all cortical layers except layer I (Fig. 3a1,b1), which nevertheless showed diffuse GFP labelling presumably originating from the ascending axon of Martinotti cells branching in this layer (DeFelipe et al., 2013). The average density of GFP-positive cells was comparable between sections immunostained for NOS-1 (49.9 ± 2.7 cells/mm^2^, n=3 mice, Fig. 3a1) and NPY (41.0 ± 7.1 cells/mm^2^, n= 3 mice, Fig. 3b1, t(2.55619)= 1.16763, p= 0.3403). A similar density of neurons immunolabeled for NOS-1 was observed in GFP positive (2.0 ± 0.1 cells/mm^2^) and negative cells (1.9 ± 0.4 cells/mm^2^, t(2.28582)= 0.215387, p= 0.8472, Fig. 3a3-4). However, the intensity of NOS-1 immunostaining was stronger in GFP-positive cells than in negative cells (Fig. 3a1,2), indicating that they likely correspond to type 1 and type 2 NO-producing interneurons, respectively. Consistent with previous reports in Sst interneurons, (Kubota et al., 1994, 2011; Karagiannis et al., 2009; Perrenoud et al., 2012a; Tricoire et al., 2013), putative type 1 NO interneurons represented only a minority of GFP-positive cells (4.0 ± 0.1%). NPY immunostaining showed a relatively high density of positive cells (29.4 ± 6.0 cells/mm^2^) and fiber plexus in superficial and deep layers (Fig. 3b2) as reported in other mammals (Hendry et al., 1984). The vast majority of NPY immunoreactive neurons were also GFP positive (81.4 ± 2.1%, Fig. 3b3-4) with the remaining NPY interneurons likely corresponding to neurogliaform cells (Cauli et al., 2004; Price et al., 2005; Karagiannis et al., 2009; Kubota et al., 2011; Perrenoud et al., 2012a), as suggested by their round immunolabeled somata (Fig. 3b3-4). More precisely, the proportion of GFP-positive cells immunostained for NPY was much higher (58.9 + 8.2%, t(2.00020)=6.70695, p= 0.0215) than those immunolabeled for NOS-1, which is known to represent a subpopulation of NPY interneurons (Estrada and DeFelipe, 1998; Karagiannis et al., 2009; Tasic et al., 2016). These observations collectively indicate that the Cre-dependent expression of the transgenes in Sst-IRES-Cre mice occurs in at least two subpopulations of interneurons that express NOS-1 and/or NPY to a greater extent.

**Figure 3:**
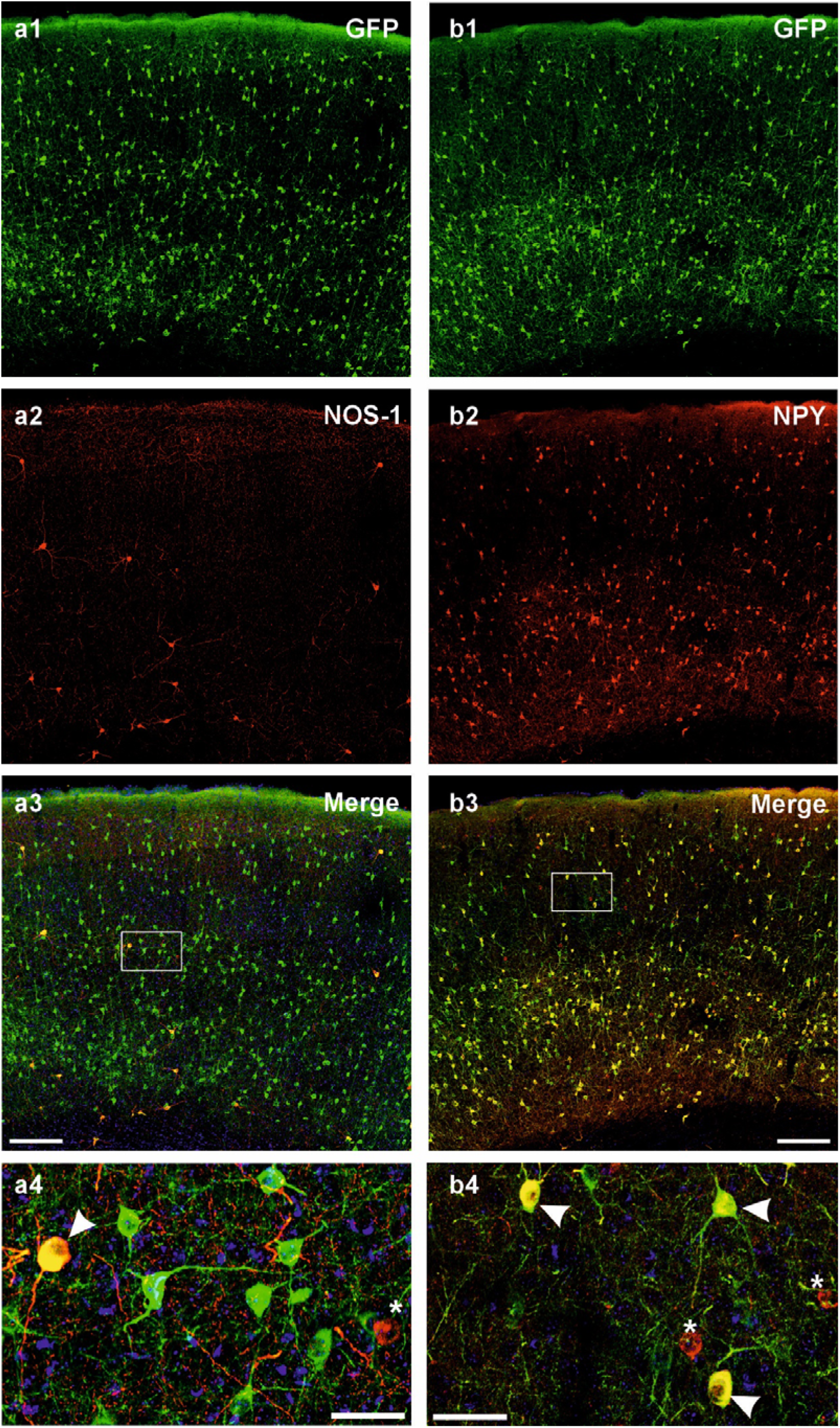
Expression of NOS-1 and NPY in Sst-Cre neurons. **(a1,b1)** Representative maximum intensity projections of 30 µm thick stacked confocal images showing GFP (green) expression in cortical slices from Sst-GCaMP6f mice. (**a2**) NOS-1 immunolabeling (red) is present in rare neurons. (**b2**) Numerous GFP-positive neurons are immunolabeled for NPY (red). (**a3,b3**) Double fluorescence images showing the co-expression of NOS-1 in rare GFP-positive neurons (a3) and NPY in numerous GFP positive neurons (b3). The white rectangles in **a3** and **b3** correspond to the zoomed regions shown in **a4** and **b4**, respectively. Arrowheads indicate GPF-positive neurons immunolabelled for NOS-1 (**a4**) or NPY (**b4)**. Asterisks denote GFP-negative neurons immunostained for NOS-1 (**a4**) or NPY (**b4**). Scale bars: 200 µm (**a3, b3**) and 40 µm (**a4,b4**).

### Vasodilation induced by 2 Hz photostimulation of Sst-ChR2 interneurons requires AP firing and NO signaling

Since a subpopulation of Sst interneurons (Fig. 3a3-4 and supplementary 3) can produce the vasodilator NO (Cauli et al., 2004), we next investigated the mechanisms by which they induce vasodilation when photostimulated at 2 Hz. We first determined whether spiking activity was required by blocking APs with the voltage-activated sodium channel blocker tetrodotoxin (TTX 1 µM (Zonta et al., 2003), n= 10 arterioles). Bath application of TTX did not change arteriolar diameter (diameter before: 17.0 ± 1.8 µm *vs.* after TTX: 16.9 ± 1.7 µm, t_(9)_= 1.0031, p=0.3420, Supplementary Fig. 4a,b), indicating that basal network activity does not influence the resting vascular tone. By contrast, it dramatically reduced the overall vascular response (-19.6 ± 10.5 %.min, p= 0.04966, Fig. 4a and d1) and the magnitude of evoked vasodilation (6.8 ± 2.4 %.min, p= 0.00844, Fig. 4d2) without altering the maximal dilation (2.2 ± 0.6%, p=0.11114, Fig. 4d3) or its time to peak (1256 ± 192 s, p= 0.07716, Fig. 4e2). In addition, the onset and the duration of vasodilation were respectively delayed (1142 ± 210 s, p= 0.00152, Fig. 4e1) and shortened (306 ± 121 s, p= 0.01623, Fig. 4e3). Of note, the vasoconstrictive events were not altered except for their onset and time to peak (Supplementary Fig. 4g1-5). These observations indicate that AP firing is mandatory for the vasodilation induced by 2 Hz photostimulation.

**Figure 4:**
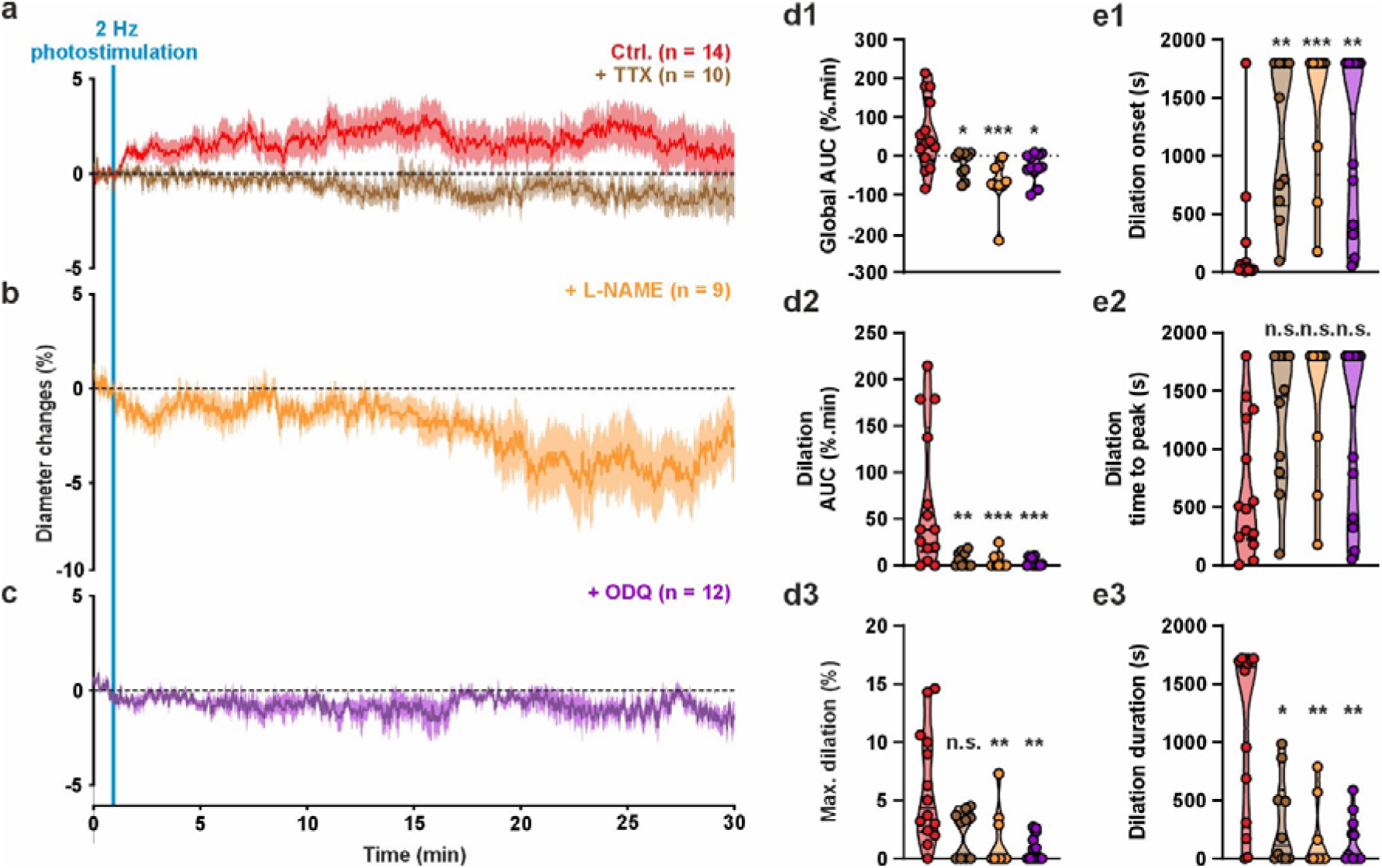
Vasodilation induced by 2 Hz photostimulation of Sst-ChR2 neurons requires spiking activity and nitric oxide signaling. **(a, b, c)** Effects of tetrodotoxin (TTX, 1 µM, brown, n= 10 arterioles from 6 mice), NOS inhibitor (L-NAME, orange, 100 µM, n= 9 arterioles from 6 mice) and ODQ (10 µM, purple, n= 12 arterioles from 7 mice), on the kinetics of the vascular response induced by 2 Hz photostimulation of Sst-ChR2 neurons. The red traces correspond to the control condition shown in Fig. 2b. The SEMs envelope the mean traces. Vasodilation is abolished by blocking action potentials with TTX **(a)**, inhibiting nitric oxide synthase with L-NAME **(b)** or sGC with ODQ **(c)**. The dashed black lines indicate the normalized initial diameters. Violin plots summarizing the effects of TTX (brown), L-NAME (orange) and ODQ (purple) on the overall vascular response **(d1**, H_(5,51)_= 16.105, p= 0.00288), magnitude **(d2**, H_(5,51)_= 18.625, p= 0.00093), maximum **(d3**, H_(5,51)_= 14.834, p=0.00506), onset **(e1**, H_(5,51)_= 20.592, p= 0.00038), time to peak **(e2**, H_(5,51)_= 8.299, p= 0.08123), and duration **(e3**, H_(5,51)_= 16.867, p= 0.00205) of vasodilation induced by 2 Hz photostimulation (red). Data are presented as individual values. Solid and dashed black bars correspond to median and quartile values. * and ** statistically different from control condition with p<0.05 and 0.01 respectively.

We next asked whether NO signaling could be responsible for the vasodilation induced by 2 Hz photostimulation. Inhibiting nitric oxide synthases with L-NAME (100 µM (Zonta et al., 2003), n = 9 arterioles) did not alter the arteriolar diameter (diameter before: 15.3 ± 2.1 µm *vs.* after L-NAME: 15.4 ± 2.3 µm, W_(9)_= -2, p=0.9453, Supplementary Fig. 4c,d) indicating that basal NO synthase activity does not influence resting vascular tone. By contrast L-NAME treatment completely abolished the evoked vasodilation as evidenced by its dramatic impact on the overall vascular response (AUC: -71.0 ± 20.4 %.min, p= 0.00042, Fig. 4b and 4d1), as well as on the magnitude (4.9 ± 2.9 %.min, p = 0.00096, Fig. 4d2) and maximal amplitude of vasodilation (1.5 ± 0.9%, p= 0.00172, Fig. 4d3). It also strongly delayed the apparent onset of vasodilation (1407 ± 210 s, p= 0.0004, Fig. 4e1) and reduced its duration (169 ± 100 s, p=0.00059, Fig. 4e3), but not the time to peak (1410 ± 210 s, H_(4,54)_= 8.299, p= 0.08123, Fig. 4e2). The overall negative responses observed under NOS inhibition suggest that vasoconstriction was uncovered. Indeed, L-NAME increased the magnitude (-70.1 ± 19.7 %.min, p= 0.00234, Supplementary Fig. 4g1) and the maximal amplitude (-8.6 ± 1.9 %, p= 0.00089, Supplementary Fig. 4g2) of vasoconstrictive events induced by 2 Hz photostimulation. It also shortened their onset (475 ± 207s, p= 0.01292, Supplementary Fig. 4g3) without affecting their time to peak (1056 ± 185 s, p= 0.06502, Supplementary Fig. 4g4) and duration (926 ± 155 s, H_(4,54)_=9.278, p= 0.05452, Supplementary Fig. 4g5). These observations suggest that the vasodilation evoked by 2 Hz photostimulation involves NO synthesis, and that its blockade unmasks vasoconstriction. To confirm that NO signaling is required, we next used ODQ (10 µM (Hepp et al., 2007), n= 12 arterioles) to inhibit the sGC, the intracellular receptor responsible for NO-mediated vasodilation. Similar to NOS inhibition, blockade of NO-signaling did not alter the basal diameter (diameter before: 19.3 ± 2.9 µm *vs.* after ODQ: 19.2 ± 2.9 µm, W_(12)_= -14, p=0.6055, Supplementary Fig. 4c,d) confirming that NO signaling does not influence resting vascular tone. Similar to NOS inhibition by L-NAME, ODQ dramatically altered the overall vascular response (-27.6 ± 10.2 %.min, p= 0.02253, Fig. 4c and d1) and reduced the magnitude (3.0 ± 1.2%, p=0.00093, Fig. 4d2) and maximal amplitude of vasodilation (0.9 ± 0.3 %, p= 0.00172, Fig. 4d3). It also strongly delayed the vasodilation onset (1119 ± 216 s, p= 0.00130, Fig. 4e1), and reduced its duration (147 ± 58 s, p= 0.00123, Fig. 4e3) without impacting the time to peak of vasodilation (1119 ± 216 s, Fig. 4e2). In contrast to L-NAME treatment, ODQ, did not alter the magnitude, amplitude, or kinetic properties of vasoconstrictive events (Supplementary Fig. 4g1-5). Taken together, these results indicate that AP firing is mandatory for the vasodilation evoked by 2 Hz photostimulation, which is mediated by NO synthesis and signaling. Furthermore, they show that 2 Hz optogenetic stimulation induces vasoconstriction that is masked by NO-dependent vasodilation.

### NPY-induced vasoconstriction involves the Y1 receptor independently of spiking activity

Since the majority of Sst interneurons are positive for NPY (Fig. 3b1,2) and virtually all cortical type 1 NO-producing interneurons express this neuropeptide (Dawson et al., 1991; Estrada and DeFelipe, 1998; Kubota et al., 1994, 2011; Perrenoud et al., 2012a), NPY could be responsible for the vasoconstriction observed with 20 Hz photostimulation, but also with 2 Hz photostimulation under NOS inhibition. We first verified that NPY induced vasoconstriction in a dose-dependent manner (Supplementary Fig. 5). 10 nM NPY (n= 11 arterioles) evoked very small vasoconstriction (Supplementary Fig. 5b e1-3), that only differed form control condition (n= 3 arterioles) for their onset (10 nM: 548 ± 194 s, p= 0.04430, Supplementary Fig. 5f1). 100 nM (n= 10 arterioles) and 1 µM NPY (n= 9 arterioles) evoked much stronger vasoconstriction, as evidenced by their magnitude (100 nM: -154.9 ± 40.7 %.min, p= 0.01154; 1 µM: -162.5 ± 23.0 %.min, p= 0.00474, Supplementary Fig. 5e1), maximal amplitude (100 nM: -13.1 ± 2.2 %, p= 0.02023; 1 µM: -25.1 ± 3.4 %, p= 0.00062, Supplementary Fig. 5e2). Vasoconstriction evoked by 100 nM and 1 µM occurred also much faster (100 nM: 282 ± 43 s, p= 0.04430; 1µM: 147 ± 18 s, p= 0.00281, Supplementary Fig. 5e3) and time to peak (100 nM: 545 ± 116 s, p= 0.02912; 1µM: 311 ± 38 s, p= 0.00395, Supplementary Fig. 5e4). They also lasted longer (998 ± 124 s, p= 0.01731; 1µM: 1234 ± 104s, p= 0.00113, Supplementary Fig. 5e5). Since the vasoconstriction induced by 20 Hz optogenetic stimulation of Sst-ChR2 and 100 nM NPY were of similar magnitude and maximal amplitude (Fig. 2f2,3 and supplementary Fig. 5e1,2) we next evaluated whether the NPY Y1 antagonist BIBP3226 could abolish the vasoconstriction induced by 100 nM. Pretreatment with BIBP3226 (1µM (Perrenoud et al., 2012b), n= 10 arterioles) dramatically reduced the vasoconstriction induced by 100 nM NPY (Fig. 5a), as evidenced by its effect on the magnitude (-8.0 ± 7.6 %.min, p= 0.00181, Fig. 5c1) and maximum amplitude (-0.8 ± 0.7 %, p= 0.00144, Fig. 5c2). BIBP3226 also dramatically delayed the onset (1486 ± 212 s, p= 0.01467, Fig. 5c3) and time to peak (1622 ± 137 s, p= 0.00207, Fig. 5c4) of NPY-induced vasoconstriction and reduced its duration (175 ± 148 s, p= 0.01261, Fig. 5c5). These observations confirm that NPY induces strong arteriolar vasoconstriction via Y1 receptors. Since Y1 receptor is expressed by smooth muscle cells (Vanlandewijck et al., 2018; Zhang et al., 2024) but also by neurons (Smith et al., 2019; Tasic et al., 2016; Zeisel et al., 2015) it has been proposed that NPY-induced vasoconstriction could be mediated by intermediary neuronal types (Kaplan et al., 2020). To rule out this possibility we investigated the impact of AP firing inhibition on NPY-induced vasoconstriction. We found that TTX treatment (1 µM, n= 9 arterioles) had no effect on the magnitude (-173.7 ± 58.8 %.min, p= 0.62911, Fig. 5c1), maximum amplitude (-14.3 ± 4.2 %, p= 0.64910, Fig. 5c2), onset (554 ± 239s, p= 0.88537, Fig. 5c3), time to peak (752 ± 211 s, p= 0.45002, Fig. 5c4) and duration (903 ± 199s, p= 0.80606, Fig. 2c5) of vasoconstriction induced by 100 nM NPY. Collectively these observations indicate that NPY induces vasoconstriction via the Y1 receptor independently of neuronal activity.

**Figure 5:**
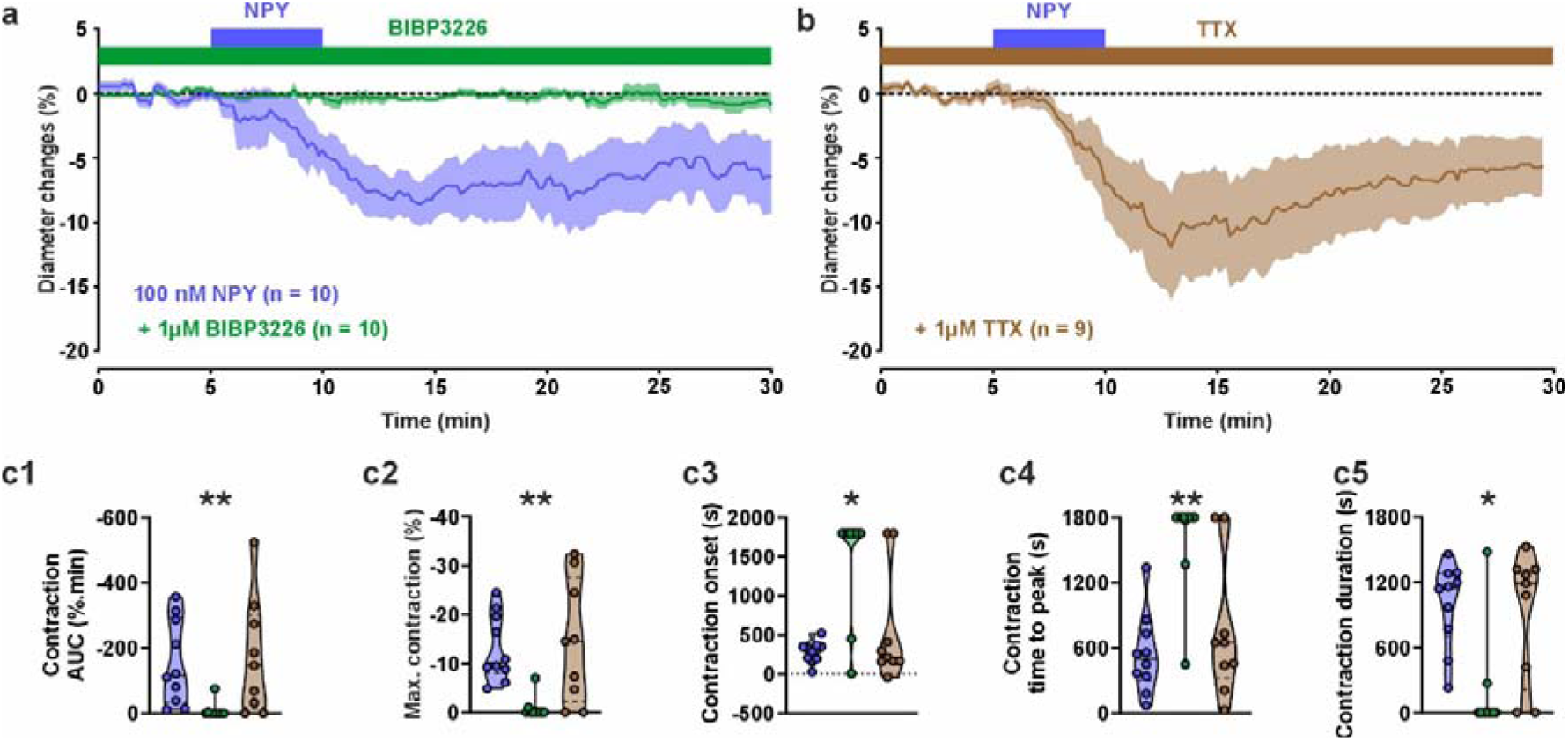
NPY-induced vasoconstriction involves the Y1 receptor independently of spiking activity. **(a,b)** Kinetics of diameter changes induced by 5 minutes exogenous application of 100 nM NPY (blue bars) under control condition (**a**, blue trace) and in presence of 1 µM BIBP3226 (**a**, green trace, n= 10 arterioles from 6 mice) or 1 µM TTX (**b**, brown trace, n= 9 arterioles from 4 mice). The SEMs envelope the mean traces. Violin plots summarizing the effects of BIBP 3226 (green) and TTX (brown) on the on the magnitude **(c1**, H_(4,32)_= 17.244, p= 0.00063), maximum **(c2**, H_(4,32)_= 17.859, p= 0.00047), onset **(c3**, H_(4,32)_= 13.529, p= 0.00362), time to peak **(c4**, H_(4,32)_= 16.209, p= 0.00103), and duration **(c5**, H_(4,32)_= 13.557, p= 0.00357) of vasoconstriction induced by NPY. The selective NPY Y1 receptor antagonist BIBP3226 but not TTX abolished the NPY-induced vasoconstriction. Data are presented as individual values. Solid and dashed black bars correspond to median and quartile values. * and ** statistically different from the 100 nM control condition with p< 0.05 and 0.01, respectively.

### Vasoconstriction induced by 20 Hz photostimulation of Sst-ChR2 interneurons requires AP firing and NPY Y1 signaling

To investigate the mechanisms responsible for the vasoconstriction induced by 20 Hz optogenetic stimulation of Sst-ChR2 interneurons, we first evaluated the requirement of AP firing. TTX treatment (1µM, n= 10 arterioles) dramatically reduced the magnitude (-14.1 ± 5.8, %.min, p= 0.01810, Fig. 6a, c2), maximum amplitude (-1.9 ± 0.5 %, p= 0.00092, Fig. 6c3) of vasoconstriction induced by 20 Hz photostimulation. It also delayed the vasoconstriction onset (990 ± 236 s, p= 0.01993, Fig. 6d1) without significantly altering its apparent time to peak (1163 ± 187 s, Fig. 6d2) and duration (571 ± 196 s, Fig. 6d3). TTX had also no impact on vasodilatory events evoked by 20 Hz photostimulation (Supplementary Fig. 6c1-5). These observations indicate that AP firing is mandatory for vasoconstriction induced by 20 Hz optogenetic stimulation of Sst-ChR2 interneurons. We next investigated whether this vascular response could be mediated by NPY. Antagonizing Y1 receptor in Sst-ChR2 slices with BIBP3226 (1µM, n=10 arterioles) did not alter the arteriolar diameter (diameter before: 17.1 ± 2.0 µm *vs.* after BIBP3226: 17.0 ± 2.0 µm, t_(9)_= 0.82945, p= 0.4283, Supplementary Fig. 6a,b) indicating that tonic NPY level does not influence resting vascular tone. In contrast, BIBP3226 dramatically reduced the magnitude (-21.8 ± 8.6 %.min, p= 0.01810, Fig. 6a,c2) and the maximum amplitude (-2.0 ± 0.6 %, p= 0.00092, Fig. 6c3) of vasoconstriction induced by 20 Hz optogenetic stimulation. It also delayed the onset of vasoconstriction (934 ± 244 s, p= 0.01993, Fig 6d1) without altering its time to peak (1148 ± 201s, Fig. 6d2) and duration (643 ± 224s, Fig. 6d3). Similar to TTX, BIBP3226 had no impact on the vasodilatory events induced by 20 Hz Photostimulation (Supplementary Fig. 6c1-5). Taken together, these observations indicate that vasoconstriction induced by 20 Hz photostimulation of Sst-ChR2 neurons requires both AP firing and NPY signaling through the Y1 receptor.

**Figure 6:**
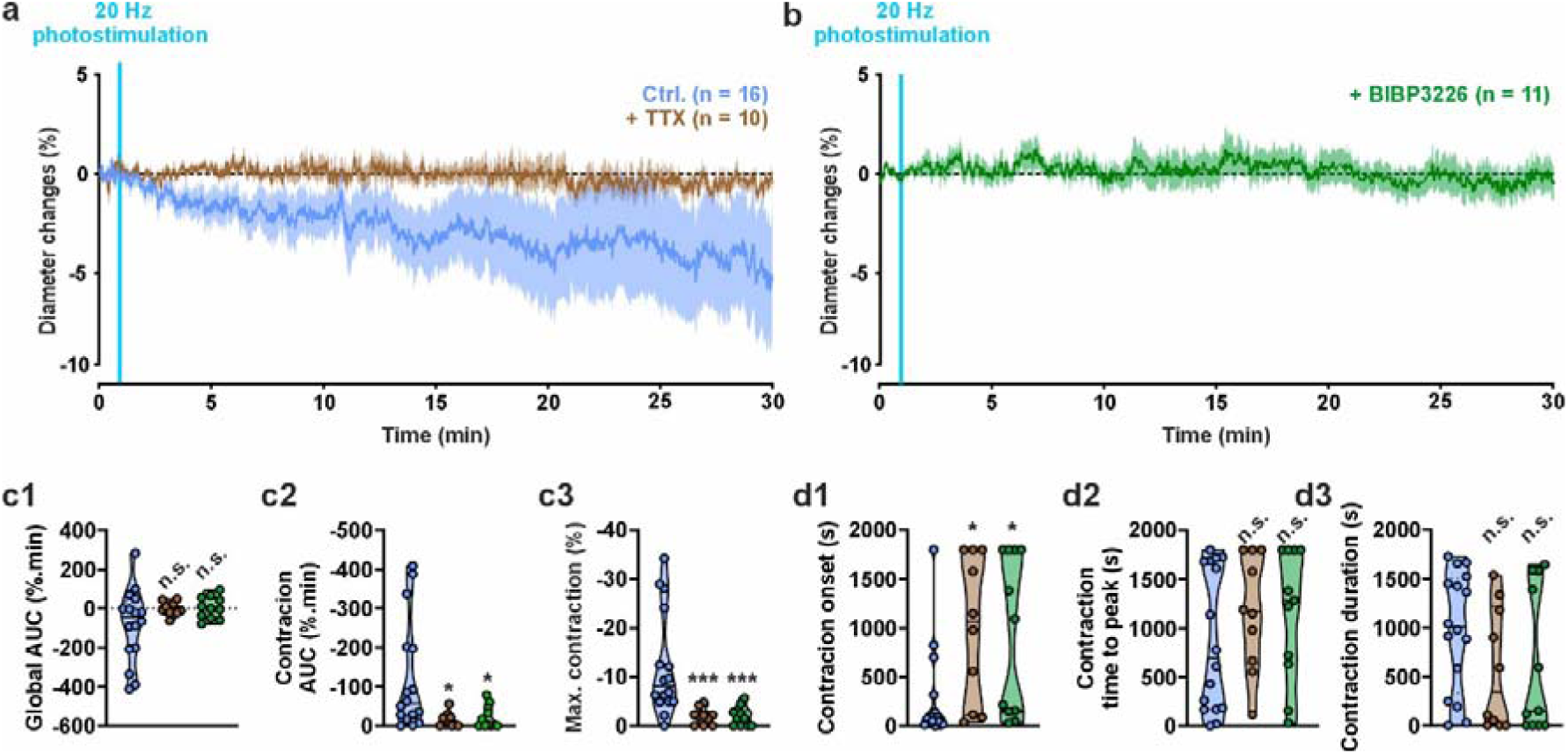
Vasoconstriction induced by 20 Hz photostimulation of Sst-ChR2 neurons requires spiking activity and NPY signaling. **(a, b)** Effects of tetrodotoxin (TTX, 1 µM, brown, n= 10 arterioles from 5 mice) and the selective Y1 receptor antagonist (BIBP 3226, 1 µM, green, n= 11 arterioles from 9 mice) on the kinetics of the vascular response induced by 20 Hz photostimulation of Sst-ChR2 neurons. The blue trace corresponds to the control condition shown in Fig. 2e. The SEMs envelope the mean traces. Vasoconstriction is abolished by blocking action potentials with TTX **(a)** or by the selective antagonist of Y1 receptor, BIBP 3226 **(b)**. Violin plots summarizing the effects of TTX (brown) and BIBP 3226 (green) on the overall vascular response **(c1,** F*_(3,22.04)_= 2.557, p= 0.08117), magnitude **(c2,** H_(4,41)_= 10.774, p= 0.01301), maximum **(c3,** H_(4,41)_= 19.152, p= 0.00025), onset **(d1,** H_(4,41)_= 10.671, p= 0.01364), time to peak **(d2,** H_(4,41)_= 6.373, p= 0.09481), and duration **(d3,** H_(4,41)_= 4.726, p= 0.19300) of vasoconstriction induced by 20 Hz photostimulation (cyan). Data are presented as individual values. Solid and dashed black bars correspond to median and quartile values. *, ** and *** statistically different from control condition with p<0.05, p<0,01 and 0.001 respectively.

## DISCUSSION

Here we used optogenetics in mouse cortical slices to show that Sst interneurons exert a bimodal vascular response that requires AP firing and whose polarity depends on the stimulation frequency. The vasodilation induced by 2 Hz photostimulation involves NO synthesis and signaling via the sGC, whereas the vasoconstriction induced by 20 Hz photostimulation implies NPY and activation of its Y1 receptor. Consistently, histochemical investigations revealed that subpopulations of photostimulated Sst interneurons express the NOS-1 and/or NPY (Fig. 7).

**Figure 7:**
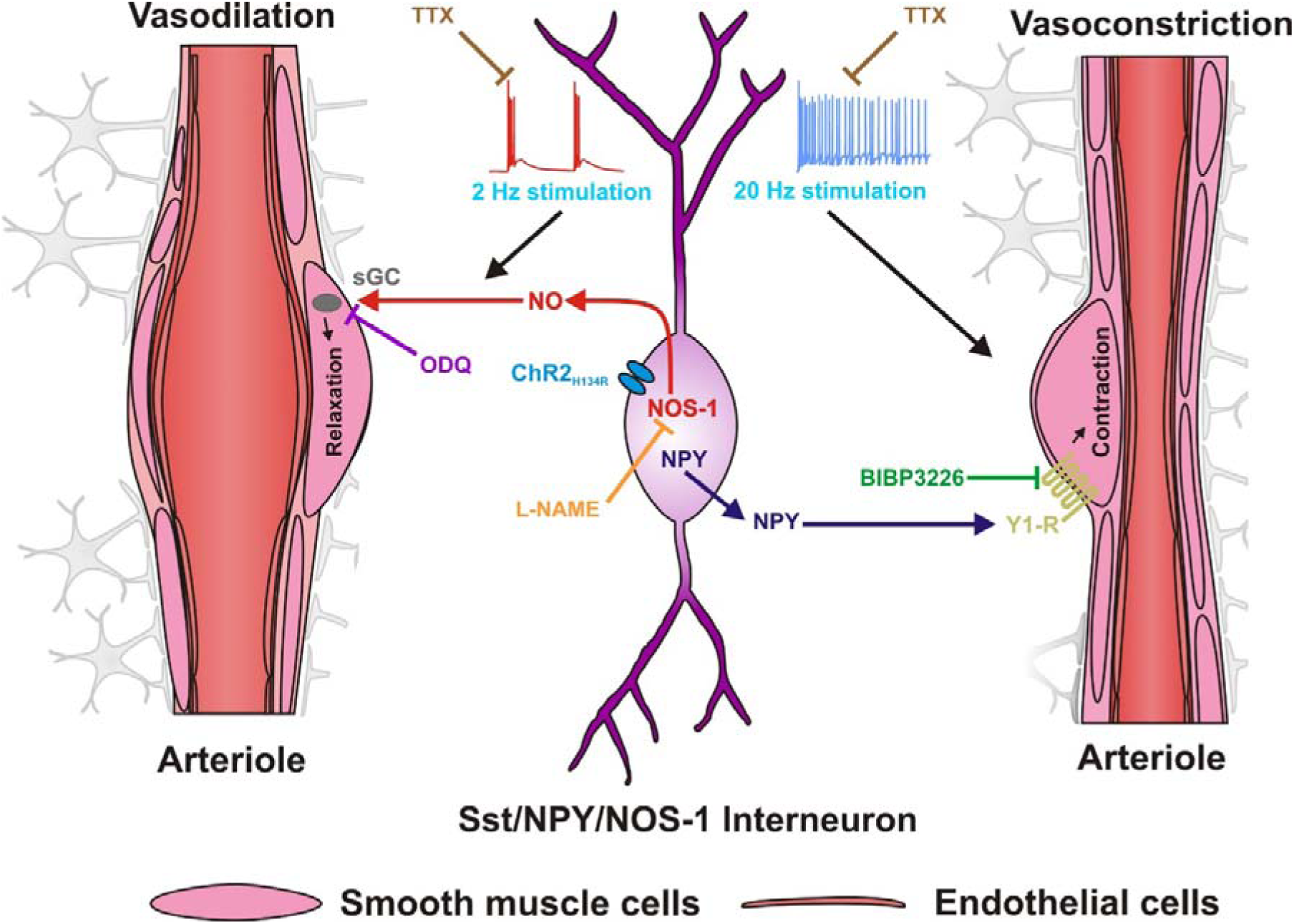
Schematic representation of the mechanisms of bidirectional control of neurovascular coupling by Sst-ChR2 interneurons. Bidirectional responses are shown on both sides of the Sst-NPY-NOS-1 interneuron. **Left:** 2 Hz stimulation of Sst-ChR2 interneurons activates NOS-1 and releases NO which in turn activates the sGC of smooth muscle cells leading to their relaxation and vasodilation. This effect is abolished by TTX (brown) and by inhibition of NOS with L-NAME (orange) and sGC with ODQ (purple). **Right**: 20 Hz photostimulation of Sst-ChR2 interneurons induces NPY release and activation of vascular receptor Y1 leading to smooth muscle cell contraction and vasoconstriction. This effect is suppressed by the TTX (brown) and the Y1 receptor antagonist, BIPB3226 (green).

### Frequency dependence of the neuronal and vascular responses

The observation that Sst interneurons respond supralinearly to low-frequency optogenetic stimulation is reminiscent of the higher impedance observed at low frequency in hippocampal Sst-positive oriens lacunosum-moleculare cells (Zemankovics et al., 2010), which depends on a high capacitance and a pronounced hyperpolarisation-activated cation current I_h_, two features shared by cortical Sst interneurons (Halabisky et al., 2006; Karagiannis et al., 2009, 2021; Perrenoud et al., 2012a). Indeed, according to Ohm’s law, a higher impedance at low frequency would have resulted in a larger voltage response induced by the ChR2 photocurrents, resulting in a higher number of spikes per light pulse. We found that depending on the AP firing frequency, Sst interneurons can induce vasodilation or vasoconstriction. These results confirm and extend our previous observations in rat cortical slices showing that the evoked firing of single Sst interneurons at a frequency of at least 8 Hz results in vasoconstriction (Cauli et al., 2004). Similarly, the *in vivo* optogenetic stimulation of GABAergic interneurons was found to induce hyperemia/vasodilation, which can be followed by vasoconstriction (Anenberg et al., 2015; Uhlirova et al., 2016). In addition, photostimulation of Sst interneurons also induced a hyperemic response, followed by a decrease in blood flow that became more pronounced with increasing duration of stimulation (Krawchuk et al., 2019; Lee et al., 2020). Our findings are further supported by the observations showing that the selective optogenetic, chemogenetic or pharmacological stimulation of NOS-1 expressing Sst interneurons induced hyperemia/vasodilation (Echagarruga et al., 2020; Ruff et al., 2024) also support our findings. Nevertheless compared to those *in vivo,* the responses we observed in slices were notably much slower and longer lasting (Krawchuk et al., 2019; Lee et al., 2020). This is likely due to the lower temperature in *ex vivo* experiments which has been reported to slowdown the synthesis of vasoactive messenger (Rancillac et al., 2006) and subsequent reactions. In contrast to *in vivo* observations (Uhlirova et al., 2016; Krawchuk et al., 2019; Lee et al., 2020), the *ex vivo* magnitude of vasoconstriction was greater than that of vasodilation. Here, the absence of vascular tone in slices, which favors vasoconstriction (Blanco et al., 2008), probably led to an underestimation of vasodilation. On the other hand the absence of blood flow in the slices may have reduced the scavenging of NO by hemoglobin (Rancillac et al., 2006), whose *in vivo* concentration locally increases during functional hyperemia (Devor et al., 2007; Lee et al., 2020), thereby prolonging *ex vivo* vasodilation.

### Cellular and molecular mechanisms of vasodilation and vasoconstriction

We found that the vasodilation induced by the 2 Hz optogenetic stimulation was dependent on AP firing, NO synthesis and activation of soluble guanylate cyclase (sGC). The mandatory role of APs indicates that Ca^2+^ entry via ChR2 alone is insufficient to drive vasodilation. This likely reflects the essential role of voltage-gated Ca^2+^ channels in somatic Ca^2+^ elevation (Mao et al., 2001) and the subsequent Ca^2+^-dependent activation of NOS-1, NO release and arteriolar vasodilation (Rancillac et al., 2006; Attwell et al., 2010; Cauli and Hamel, 2010; Mishra et al., 2016; Iadecola, 2017; Cauli and Hamel, 2018). Consistently, some supragranular NOS-1 neurons showed an increase in somatic Ca^2+^ prior to vasodilation evoked by whisker stimulation and/or spontaneous locomotion in awake head-fixed mice whereas others showed a gradual and delayed activation (Ahn et al., 2023). Although type 1 NOS-1 interneurons are preferentially located in deep layers (Perrenoud et al., 2012a), our histochemical observations revealed the presence of NOS-1 positive Sst-Cre interneurons in superficial layers. Together with the high excitability of Sst interneurons (Karagiannis et al., 2009, 2021; Perrenoud et al., 2012a), these observations suggest that some of these interneurons correspond to the fast responder NOS-1 neurons (Ahn et al., 2023). The abolition of the optogenetically induced vasodilation evoked by 2 Hz photostimulation by both NOS and sGC inhibition strongly suggests that it is mediated by NO synthesis, release and direct action on smooth muscle cells, leading to their relaxation (Fig. 7). However, NOS inhibition revealed a vasoconstriction, most likely mediated by NPY (Uhlirova et al., 2016), that was not observed when sGC or AP firing was blocked. These observations suggest that a NO-sensitive, but soluble guanylate cyclase-independent, mechanism of vasodilation counterbalanced the vasoconstriction disclosed by L-NAME. Active astrocytes, which have been shown to produce vasodilatory epoxyeicosatrienoic acids via the P450 epoxygenase (Alkayed et al., 1997), may be responsible for such a vasodilation. Indeed, astrocytes have been reported to be robustly activated by the repetitive firing of Sst interneurons (Mariotti et al., 2018), and the P450 expoygenase, which is expressed in arteriolar but not cappillary endfeet (Mishra et al., 2016), is inhibited by NO (Roman, 2002; Metea and Newman, 2006; Attwell et al., 2010).

We found that vasoconstriction predominates over vasodilation when Sst-ChR2 interneurons are photostimulated at 20 Hz. This reflects the much higher number of NPY-positive Sst-ChR2 interneurons compared to type 1 NOS-1 interneurons (Karagiannis et al., 2009; Perrenoud et al., 2012a) and the dense network of NPY positive terminals on blood vessels (Abounader and Hamel, 1997). Furthermore, the release of NPY from large dense-core vesicles requires a higher firing frequency than the release of neurotransmitters from synaptic vesicles (Baraban and Tallent, 2004). It is also possible that the release of NO requires a lower firing frequency than that of NPY. We found that optogenetically induced vasoconstriction was almost completely abolished by the antagonism of Y1 receptors, suggesting that this effect was essentially mediated by NPY, as reported for the optogenetic stimulation of GABAergic interneurons (Uhlirova et al., 2016). We cannot exclude the indirect contribution of Sst and GABA which can exert vasoconstrictive effects (Long et al., 1992; Fergus and Lee, 1997). Indeed, the repetitive firing of Sst interneurons robustly activates astrocytes via Sst and GABA_B_ receptors (Mariotti et al., 2018), which could in turn induce vasoconstriction via 20-hydroxyeicosatetraenoic acid (Mulligan and MacVicar, 2004) and/or K+ (Girouard et al., 2010). However, Sst is not as efficient as NPY in inducing vasoconstriction in cortical slices (Cauli et al., 2004), and the amount of Sst in the cerebral cortex is reduced in Sst-Cre mice (Viollet et al., 2017), but not that of NPY. On the other hand, the vasoconstrictor effect of the GABA_B_ receptor activation requires AP firing (Fergus and Lee, 1997), suggesting that it is mediated by neurons and not by astrocytes. Thus, it is more likely that the vasoconstriction induced by 20 Hz optogenetic stimulation of Sst-ChR2 interneurons is essentially mediated by NPY and the Y1 receptors.

### Physiological relevance of the bidirectional control of neurovascular coupling by Sst-interneurons

The ability of Sst interneurons to drive a bimodal vascular response *ex vivo* depending on firing frequency is consistent with *in vivo* observations showing that increasing the duration and/or the frequency of photostimulation of Sst-ChR2 interneurons favors the observation of decreased blood flow, in addition to hyperemia (Krawchuk et al., 2019; Lee et al., 2020). However, the evoked decrease in blood flow appears to be more pronounced at a site different from the site of hyperemia (Lee et al., 2020). This is reminiscent of the center/surround pattern of vasodilation/vasoconstriction observed *in vivo* in response to sensory stimulation, with the vasodilation in the activated area preceding the peripheral vasoconstriction (Devor et al., 2007; Boorman et al., 2010; Uhlirova et al., 2016). Because fast-responding NOS-1 neurons are activated before the onset of vasodilation (Ahn et al., 2023), they are most likely involved in the initiation of vasodilation. Consistently, local NO production precedes functional hyperemia but disappears before the end of stimulation (Buerk et al., 2003), presumably because of the short half-life of NO (Wood and Garthwaite, 1994) and its increased scavenging induced by functional hyperemia. The delayed and/or peripheral vasoconstriction (Devor et al., 2007; Uhlirova et al., 2016; Krawchuk et al., 2019; Lee et al., 2020) could be due to its blunting by early occurring vasodilation, a higher level of activity required for NPY release (Baraban and Tallent, 2004), as suggested by our observations, and/or a different site of action of NO and NPY (Estrada and DeFelipe, 1998).

Sst/NOS-1/NPY neurons largely correspond to long-range GABAergic neurons (Tomioka et al., 2005) and have been shown to locally induce vasodilation (Echagarruga et al., 2020; Ruff et al., 2024). It remains to determine whether their interhemispheric long-range projection (Ruff et al., 2024) could explain the vasoconstriction observed contralaterally to the activated area (Devor et al., 2008) when their activity is elevated. Sst/NOS-1/NPY also correspond to cortical sleep active neurons (Gerashchenko et al., 2008; Kilduff et al., 2011). Furthermore, Sst and Sst/NOS-1 interneurons are involved in slow and delta wave activities, respectively (Morairty et al., 2013; Funk et al., 2017; Zielinski et al., 2019), a frequency range associated with sleep that we showed was the most efficient at activating them. Interestingly, oscillating vasodilation of large amplitude and a vasoconstriction are observed during non REM sleep and upon awakening, respectively (Turner et al., 2020; Bojarskaite et al., 2023). Whether the coordinated action of NO and NPY could participate in these hemodynamic changes in a frequency-dependent manner remains also to be determined.

### Bidirectional control of neurovascular coupling by Sst interneurons in pathological conditions

Several neurological disorders, such as epilepsy (Sada et al., 2015; Gaxiola-Valdez et al., 2017) or Alzheimer’s disease (Palop et al., 2007; Kimbrough et al., 2015; Le Douce J. et al., 2020), associate hyperactivity with gliovascular and metabolic deficits that are likely to alter the control of neurovascular coupling by Sst interneurons. For example, cerebral lactate, which is produced by astrocytes from glucose during excitatory transmission (Pellerin and Magistretti, 1994; Cauli et al., 2023), is dramatically increased in epilepsy (Slais et al., 2008). Drugs that interfere with lactate metabolism by neurons have emerged as a promising strategy for treating epilepsy (Sada et al., 2015), an effect that involves the neuronal uptake of lactate, its oxidative metabolism and the subsequent closure of ATP-sensitive potassium channels. Since Sst interneurons functionally express these channels (Karagiannis et al., 2021) and are among the neurons showing the highest mitochondrial content (Gulyas et al., 2006), their activity is likely to be efficiently enhanced by the presence of lactate and its oxidative metabolism. This could favor the GABAergic inhibition of pyramidal cells (Silberberg and Markram, 2007), but also decreased glutamatergic transmission (Bacci et al., 2002), and Y1-mediated vasoconstriction (Uhlirova et al., 2016), thereby mitigating lactate-induced hyperactivity (Sada et al., 2015).

Hyperactivity has also been reported in several models of Alzheimer’s disease (Palop et al., 2007; Bezzina et al., 2015; Zott et al., 2019), and loss of Sst interneurons has been observed in patients (Gaspar et al., 1989) and in the early stages of a mouse model of the disease (Ramos et al., 2006). Furthermore, Sst/NOS-1/NPY interneurons play a key role in non-REM sleep (Gerashchenko et al., 2008; Morairty et al., 2013; Zielinski et al., 2019), during which the fluctuations in the perivascular space favor the clearance of β-amyloid peptides (Iliff et al., 2013; Xie et al., 2013; Bojarskaite et al., 2023). These observations suggest that the control of NVC by Sst interneurons may be altered early, accelerating amyloid plaque accumulation and disease progression. However, NVC has been shown to be impaired (Niwa et al., 2002; Kimbrough et al., 2015; Lourenco et al., 2017), unchanged (Sharp et al., 2020) or even enhanced (Shabir et al., 2020; Kim et al., 2024), making it difficult to assess its alterations. Whether these differences depend on the mouse model, the stage of disease progression, or the experimental condition remains to be investigated.

## Conclusion

Here, using a combination of approaches, we describe the mechanisms by which Sst interneurons bidirectionally control NVC as a function of their firing frequency. Vasodilation was preferentially evoked at low-frequency photostimulation and primarily involves NO release and activation of its vascular receptor sGC. In contrast, vasoconstriction was more reliably induced at high frequency and involves NPY release and activation of the vascular receptor Y1. This finding will help to update the interpretation of the functional brain imaging signals used to map network activity in health and disease (Iadecola, 2017; Zhang and Raichle, 2010). This bimodal control of neurovascular coupling may also help to define therapeutic targets for epilepsy and Alzheimer’s disease in which hyperactivity, gliovascular and metabolic deficits overlap (Palop and Mucke, 2010; Sada et al., 2015; Gaxiola-Valdez et al., 2017).

## Data availability statement

The data that support the results of this study are available from the corresponding author upon reasonable request.

## Acknowledgements

We acknowledge the invaluable support of the animal (RongIBPS) and imaging facilities of the IBPS. Financial support was provided by grants from the Agence Nationale de la Recherche (ANR-17-CE37-0010-03, B.C.; ANR-20-CE14-0025, D.L.; ANR-23-CE14-0038-01, B.C.), and the i-Bio initiative of Sorbonne University (B.C.). B.M was supported by a fellowship form Centre National de la Recherche Scientifique. B.L.G. and E.B. were supported by fellowships from Fondation pour la Recherche sur Alzheimer.

## Author contributions

B.M., J.O.R, S.P. B.L.G., M.T., and E.B. designed experiments, acquired and analyzed the data, and edited the manuscript. B.M. and B.C. drafted the manuscript. L.R.V.C. edited the manuscript. I.D. designed experiments and edited the manuscript. H.S. analyzed the data. D.L. provided resources and edited the manuscript. B.C. designed experiments, analyzed the data and edited the manuscript.

## REFERENCES

Abounader R, Hamel E. 1997. Associations between neuropeptide Y nerve terminals and intraparenchymal microvessels in rat and human cerebral cortex. JComp Neurol 388:444–453.

Ahn SJ, Anfray A, Anrather J, Iadecola C. 2023. Calcium transients in nNOS neurons underlie distinct phases of the neurovascular response to barrel cortex activation in awake mice. J Cereb Blood Flow Metab 43:1633–1647. doi:10.1177/0271678X231173175

Alkayed NJ, Birks EK, Narayanan J, Petrie KA, Kohler-Cabot AE, Harder DR. 1997. Role of P-450 arachidonic acid epoxygenase in the response of cerebral blood flow to glutamate in rats. Stroke 28:1066–1072.

Anenberg E, Chan AW, Xie Y, LeDue JM, Murphy TH. 2015. Optogenetic stimulation of GABA neurons can decrease local neuronal activity while increasing cortical blood flow. JCerebBlood Flow Metab.

Ascoli GA, Alonso-Nanclares L, Anderson SA, Barrionuevo G, avides-Piccione R, Burkhalter A, Buzsaki G, Cauli B, DeFelipe J, Fairen A, Feldmeyer D, Fishell G, Fregnac Y, Freund TF, Gardner D, Gardner EP, Goldberg JH, Helmstaedter M, Hestrin S, Karube F, Kisvarday ZF, Lambolez B, Lewis DA, Marin O, Markram H, Munoz A, Packer A, Petersen CC, Rockland KS, Rossier J, Rudy B, Somogyi P, Staiger JF, Tamas G, Thomson AM, Toledo-Rodriguez M, Wang Y, West DC, Yuste R. 2008. Petilla terminology: nomenclature of features of GABAergic interneurons of the cerebral cortex. NatRevNeurosci 9:557–568.

Attwell D, Buchan AM, Charpak S, Lauritzen M, MacVicar BA, Newman EA. 2010. Glial and neuronal control of brain blood flow. Nature 468:232–243.

Bacci A, Huguenard JR, Prince D a. 2002. Differential modulation of synaptic transmission by neuropeptide Y in rat neocortical neurons. Proc Natl Acad Sci U S A 99:17125–30. doi:10.1073/pnas.012481899

Baraban SC, Tallent MK. 2004. Interneuron Diversity series: Interneuronal neuropeptides--endogenous regulators of neuronal excitability. Trends Neurosci 27:135–142.

Benjamini Y, Hochberg Y. 1995. Controlling the False Discovery Rate: A Practical and Powerful Approach to Multiple Testing. J R Stat Soc Ser B Stat Methodol 57:289–300. doi:10.1111/j.2517-6161.1995.tb02031.x

Bezzina C, Verret L, Juan C, Remaud J, Halley H, Rampon C, Dahan L. 2015. Early Onset of Hypersynchronous Network Activity and Expression of a Marker of Chronic Seizures in the Tg2576 Mouse Model of Alzheimer’s Disease. PLOS ONE 10:e0119910. doi:10.1371/journal.pone.0119910

Blanco VM, Stern JE, Filosa JA. 2008. Tone-dependent vascular responses to astrocyte-derived signals. AmJPhysiol Heart CircPhysiol 294:H2855–H2863.

Bojarskaite L, Vallet A, Bjørnstad DM, Gullestad Binder KM, Cunen C, Heuser K, Kuchta M, Mardal K- A, Enger R. 2023. Sleep cycle-dependent vascular dynamics in male mice and the predicted effects on perivascular cerebrospinal fluid flow and solute transport. Nat Commun 14:953. doi:10.1038/s41467-023-36643-5

Boorman L, Kennerley AJ, Johnston D, Jones M, Zheng Y, Redgrave P, Berwick J. 2010. Negative blood oxygen level dependence in the rat: a model for investigating the role of suppression in neurovascular coupling. JNeurosci 30:4285–4294.

Buerk DG, Ances BM, Greenberg JH, Detre JA. 2003. Temporal dynamics of brain tissue nitric oxide during functional forepaw stimulation in rats. Neuroimage 18:1–9.

Cauli B, Audinat E, Lambolez B, Angulo MC, Ropert N, Tsuzuki K, Hestrin S, Rossier J. 1997. Molecular and physiological diversity of cortical nonpyramidal cells. JNeurosci 17:3894–3906.

Cauli B, Dusart I, Li D. 2023. Lactate as a determinant of neuronal excitability, neuroenergetics and beyond. Neurobiol Dis 184:106207. doi:10.1016/j.nbd.2023.106207

Cauli B, Hamel E. 2018. Brain Perfusion and Astrocytes. Trends Neurosci 41:409–413.

Cauli B, Hamel E. 2010. Revisiting the role of neurons in neurovascular coupling. Front Neuroenergetics 2:9.

Cauli B, Tong XK, Rancillac A, Serluca N, Lambolez B, Rossier J, Hamel E. 2004. Cortical GABA interneurons in neurovascular coupling: relays for subcortical vasoactive pathways. JNeurosci 24:8940–8949.

Choi M, Yoon J, Choi C. 2010. Label-free optical control of arterial contraction. JBiomedOpt 15:015006.

Dawson TM, Bredt DS, Fotuhi M, Hwang PM, Snyder SH. 1991. Nitric oxide synthase and neuronal NADPH diaphorase are identical in brain and peripheral tissues. ProcNatlAcadSciUSA 88:7797–7801.

DeFelipe J, Lopez-Cruz PL, avides-Piccione R, Bielza C, Larranaga P, Anderson S, Burkhalter A, Cauli B, Fairen A, Feldmeyer D, Fishell G, Fitzpatrick D, Freund TF, Gonzalez-Burgos G, Hestrin S, Hill S, Hof PR, Huang J, Jones EG, Kawaguchi Y, Kisvarday Z, Kubota Y, Lewis DA, Marin O, Markram H, McBain CJ, Meyer HS, Monyer H, Nelson SB, Rockland K, Rossier J, Rubenstein JL, Rudy B, Scanziani M, Shepherd GM, Sherwood CC, Staiger JF, Tamas G, Thomson A, Wang Y, Yuste R, Ascoli GA. 2013. New insights into the classification and nomenclature of cortical GABAergic interneurons. NatRevNeurosci. doi:10.1038/nrn3444

Devienne G, Le Gac B, Piquet J, Cauli B. 2018. Single Cell Multiplex Reverse Transcription Polymerase Chain Reaction After Patch-clamp. JVisExp.

Devor A, Hillman EM, Tian P, Waeber C, Teng IC, Ruvinskaya L, Shalinsky MH, Zhu H, Haslinger RH, Narayanan SN, Ulbert I, Dunn AK, Lo EH, Rosen BR, Dale AM, Kleinfeld D, Boas DA. 2008. Stimulus-induced changes in blood flow and 2-deoxyglucose uptake dissociate in ipsilateral somatosensory cortex. JNeurosci 28:14347–14357.

Devor A, Tian P, Nishimura N, Teng IC, Hillman EM, Narayanan SN, Ulbert I, Boas DA, Kleinfeld D, Dale AM. 2007. Suppressed neuronal activity and concurrent arteriolar vasoconstriction may explain negative blood oxygenation level-dependent signal. JNeurosci 27:4452–4459.

Dodt HU, Zieglgansberger W. 1998. Visualization of neuronal form and function in brain slices by infrared videomicroscopy. HistochemJ 30:141–152.

Drew PJ. 2022. Neurovascular coupling: motive unknown. Trends Neurosci 45:809–819.

Echagarruga CT, Gheres KW, Norwood JN, Drew PJ. 2020. nNOS-expressing interneurons control basal and behaviorally evoked arterial dilation in somatosensory cortex of mice. Elife 9.

Estrada C, DeFelipe J. 1998. Nitric oxide-producing neurons in the neocortex: morphological and functional relationship with intraparenchymal microvasculature. CerebCortex 8:193–203.

Fergus A, Lee KS. 1997. GABAergic regulation of cerebral microvascular tone in the rat. JCerebBlood Flow Metab 17:992–1003.

Funk CM, Peelman K, Bellesi M, Marshall W, Cirelli C, Tononi G. 2017. Role of Somatostatin-Positive Cortical Interneurons in the Generation of Sleep Slow Waves. JNeurosci 37:9132–9148.

Gaspar P, Duyckaerts C, Febvret A, Benoit R, Beck B, Berger B. 1989. Subpopulations of somatostatin 28-immunoreactive neurons display different vulnerability in senile dementia of the Alzheimer type. Brain Res 490:1–13.

Gaxiola-Valdez I, Singh S, Perera T, Sandy S, Li E, Federico P. 2017. Seizure onset zone localization using postictal hypoperfusion detected by arterial spin labelling MRI. Brain 140:2895–2911.

Gerashchenko D, Wisor JP, Burns D, Reh RK, Shiromani PJ, Sakurai T, de la I, Kilduff TS. 2008. Identification of a population of sleep-active cerebral cortex neurons. ProcNatlAcadSciUSA.

Girouard H, Bonev AD, Hannah RM, Meredith A, Aldrich RW, Nelson MT. 2010. Astrocytic endfoot Ca2+ and BK channels determine both arteriolar dilation and constriction. ProcNatlAcadSciUSA 107:3811–3816.

Gulyas AI, Buzsaki G, Freund TF, Hirase H. 2006. Populations of hippocampal inhibitory neurons express different levels of cytochrome c. EurJNeurosci 23:2581–2594.

Halabisky BE, Shen F, Huguenard JR, Prince DA. 2006. Electrophysiological Classification of Somatostatin-positive Interneurons in Mouse Sensorimotor Cortex. JNeurophysiol.

Hartmann DA, Berthiaume AA, Grant RI, Harrill SA, Koski T, Tieu T, McDowell KP, Faino AV, Kelly AL, Shih AY. 2021. Brain capillary pericytes exert a substantial but slow influence on blood flow. NatNeurosci 24:633–645.

Hendry SH, Jones EG, Emson PC. 1984. Morphology, distribution, and synaptic relations of somatostatin- and neuropeptide Y-immunoreactive neurons in rat and monkey neocortex. JNeurosci 4:2497–2517.

Hepp R, Tricoire L, Hu E, Gervasi N, Paupardin-Tritsch D, Lambolez B, Vincent P. 2007. Phosphodiesterase type 2 and the homeostasis of cyclic GMP in living thalamic neurons. JNeurochem 102:1875–1886.

Hill RA, Tong L, Yuan P, Murikinati S, Gupta S, Grutzendler J. 2015. Regional Blood Flow in the Normal and Ischemic Brain Is Controlled by Arteriolar Smooth Muscle Cell Contractility and Not by Capillary Pericytes. Neuron 87:95–110.

Hu H, Cavendish JZ, Agmon A. 2013. Not all that glitters is gold: off-target recombination in the somatostatin-IRES-Cre mouse line labels a subset of fast-spiking interneurons. Front Neural Circuits 7:195.

Iadecola C. 2017. The Neurovascular Unit Coming of Age: A Journey through Neurovascular Coupling in Health and Disease. Neuron 96:17–42.

Iliff JJ, Wang M, Zeppenfeld DM, Venkataraman A, Plog BA, Liao Y, Deane R, Nedergaard M. 2013. Cerebral Arterial Pulsation Drives Paravascular CSF-Interstitial Fluid Exchange in the Murine Brain. JNeurosci 33:18190–18199.

Kaplan L, Chow BW, Gu C. 2020. Neuronal regulation of the blood-brain barrier and neurovascular coupling. NatRevNeurosci 21:416–432.

Karagiannis A, Gallopin T, David C, Battaglia D, Geoffroy H, Rossier J, Hillman EM, Staiger JF, Cauli B. 2009. Classification of NPY-expressing neocortical interneurons. JNeurosci 29:3642–3659.

Karagiannis A, Gallopin T, Lacroix A, Plaisier F, Piquet J, Geoffroy H, Hepp R, Naude J, Le GB, Egger R, Lambolez B, Li D, Rossier J, Staiger JF, Imamura H, Seino S, Roeper J, Cauli B. 2021. Lactate is an energy substrate for rodent cortical neurons and enhances their firing activity. Elife 10:e71424.

Kilduff TS, Cauli B, Gerashchenko D. 2011. Activation of cortical interneurons during sleep: an anatomical link to homeostatic sleep regulation? Trends Neurosci 34:10–19.

Kim TA, Cruz G, Syty MD, Wang F, Wang X, Duan A, Halterman M, Xiong Q, Palop JJ, Ge S. 2024. Neural circuit mechanisms underlying aberrantly prolonged functional hyperemia in young Alzheimer’s disease mice. Mol Psychiatry. doi:10.1038/s41380-024-02680-9

Kimbrough IF, Robel S, Roberson ED, Sontheimer H. 2015. Vascular amyloidosis impairs the gliovascular unit in a mouse model of Alzheimer’s disease. Brain 138:3716–3733.

Krawchuk MB, Ruff CF, Yang X, Ross SE, Vazquez AL. 2019. Optogenetic assessment of VIP, PV, SOM and NOS inhibitory neuron activity and cerebral blood flow regulation in mouse somato-sensory cortex 1. JCerebBlood Flow Metab 271678X19870105.

Krogsgaard A, Sperling L, Dahlqvist M, Thomsen K, Vydmantaite G, Li F, Thunemann M, Lauritzen M, Lind BL. 2023. PV interneurons evoke astrocytic Ca(2+) responses in awake mice, which contributes to neurovascular coupling. Glia.

Kubota Y, Hattori R, Yui Y. 1994. Three distinct subpopulations of GABAergic neurons in rat frontal agranular cortex. Brain Res 649:159–173.

Kubota Y, Shigematsu N, Karube F, Sekigawa A, Kato S, Yamaguchi N, Hirai Y, Morishima M, Kawaguchi Y. 2011. Selective coexpression of multiple chemical markers defines discrete populations of neocortical GABAergic neurons. CerebCortex 21:1803–1817.

Lacroix A, Toussay X, Anenberg E, Lecrux C, Ferreirós N, Karagiannis A, Plaisier F, Chausson P, Jarlier F, Burgess SA, Hillman EMC, Tegeder I, Murphy TH, Hamel E, Cauli B. 2015. COX-2-Derived Prostaglandin E2 Produced by Pyramidal Neurons Contributes to Neurovascular Coupling in the Rodent Cerebral Cortex. J Neurosci Off J Soc Neurosci 35:11791–11810. doi:10.1523/JNEUROSCI.0651-15.2015

Le Douce J., Maugard M, Veran J, Matos M, Jego P, Vigneron PA, Faivre E, Toussay X, Vandenberghe M, Balbastre Y, Piquet J, Guiot E, Tran NT, Taverna M, Marinesco S, Koyanagi A, Furuya S, Gaudin-Guerif M, Goutal S, Ghettas A, Pruvost A, Bemelmans AP, Gaillard MC, Cambon K, Stimmer L, Sazdovitch V, Duyckaerts C, Knott G, Herard AS, Delzescaux T, Hantraye P, Brouillet E, Cauli B, Oliet SHR, Panatier A, Bonvento G. 2020. Impairment of Glycolysis-Derived l-Serine Production in Astrocytes Contributes to Cognitive Deficits in Alzheimer’s Disease. Cell Metab 31:503–517.

Lee L, Boorman L, Glendenning E, Christmas C, Sharp P, Redgrave P, Shabir O, Bracci E, Berwick J, Howarth C. 2020. Key Aspects of Neurovascular Control Mediated by Specific Populations of Inhibitory Cortical Interneurons. CerebCortex 30:2452–2464.

Lee S, Hjerling-Leffler J, Zagha E, Fishell G, Rudy B. 2010. The largest group of superficial neocortical GABAergic interneurons expresses ionotropic serotonin receptors. JNeurosci 30:16796–16808.

Lin JY, Lin MZ, Steinbach P, Tsien RY. 2009. Characterization of engineered channelrhodopsin variants with improved properties and kinetics. BiophysJ 96:1803–1814.

Long JB, Rigamonti DD, Dosaka K, Kraimer JM, Martinez-Arizala A. 1992. Somatostatin causes vasoconstriction, reduces blood flow and increases vascular permeability in the rat central nervous system. JPharmacolExpTher 260:1425–1432.

Lourenco CF, Ledo A, Barbosa RM, Laranjinha J. 2017. Neurovascular uncoupling in the triple transgenic model of Alzheimer’s disease: Impaired cerebral blood flow response to neuronal-derived nitric oxide signaling. ExpNeurol 291:36–43.

Lovick TA, Brown LA, Key BJ. 1999. Neurovascular relationships in hippocampal slices: physiological and anatomical studies of mechanisms underlying flow-metabolism coupling in intraparenchymal microvessels. Neuroscience 92:47–60.

Madisen L, Garner AR, Shimaoka D, Chuong AS, Klapoetke NC, Li L, van der BA, Niino Y, Egolf L, Monetti C, Gu H, Mills M, Cheng A, Tasic B, Nguyen TN, Sunkin SM, Benucci A, Nagy A, Miyawaki A, Helmchen F, Empson RM, Knopfel T, Boyden ES, Reid RC, Carandini M, Zeng H. 2015. Transgenic mice for intersectional targeting of neural sensors and effectors with high specificity and performance. Neuron 85:942–958.

Madisen L, Mao T, Koch H, Zhuo JM, Berenyi A, Fujisawa S, Hsu YW, Garcia AJ III, Gu X, Zanella S, Kidney J, Gu H, Mao Y, Hooks BM, Boyden ES, Buzsaki G, Ramirez JM, Jones AR, Svoboda K, Han X, Turner EE, Zeng H. 2012. A toolbox of Cre-dependent optogenetic transgenic mice for light-induced activation and silencing. NatNeurosci 15:793–802. doi:10.1038/nn.3078

Mao BQ, Hamzei-Sichani F, Aronov D, Froemke RC, Yuste R. 2001. Dynamics of spontaneous activity in neocortical slices. Neuron 32:883–898.

Mariotti L, Losi G, Lia A, Melone M, Chiavegato A, Gomez-Gonzalo M, Sessolo M, Bovetti S, Forli A, Zonta M, Requie LM, Marcon I, Pugliese A, Viollet C, Bettler B, Fellin T, Conti F, Carmignoto G. 2018. Interneuron-specific signaling evokes distinctive somatostatin-mediated responses in adult cortical astrocytes. NatCommun 9:82.

Metea MR, Newman EA. 2006. Glial cells dilate and constrict blood vessels: a mechanism of neurovascular coupling. JNeurosci 26:2862–2870.

Mishra A, Reynolds JP, Chen Y, Gourine AV, Rusakov DA, Attwell D. 2016. Astrocytes mediate neurovascular signaling to capillary pericytes but not to arterioles. NatNeurosci 19:1619–1627.

Morairty SR, Dittrich L, Pasumarthi RK, Valladao D, Heiss JE, Gerashchenko D, Kilduff TS. 2013. A role for cortical nNOS/NK1 neurons in coupling homeostatic sleep drive to EEG slow wave activity. Proc Natl Acad Sci 110:20272–20277. doi:10.1073/pnas.1314762110

Mulligan SJ, MacVicar BA. 2004. Calcium transients in astrocyte endfeet cause cerebrovascular constrictions. Nature 431:195–199.

Niwa K, Kazama K, Younkin SG, Carlson GA, Iadecola C. 2002. Alterations in cerebral blood flow and glucose utilization in mice overexpressing the amyloid precursor protein. NeurobiolDis 9:61–68.

Palop JJ, Chin J, Roberson ED, Wang J, Thwin MT, Bien-Ly N, Yoo J, Ho KO, Yu GQ, Kreitzer A, Finkbeiner S, Noebels JL, Mucke L. 2007. Aberrant excitatory neuronal activity and compensatory remodeling of inhibitory hippocampal circuits in mouse models of Alzheimer’s disease. Neuron 55:697–711.

Palop JJ, Mucke L. 2010. Amyloid-beta-induced neuronal dysfunction in Alzheimer’s disease: from synapses toward neural networks. NatNeurosci 13:812–818.

Pellerin L, Magistretti PJ. 1994. Glutamate uptake into astrocytes stimulates aerobic glycolysis: a mechanism coupling neuronal activity to glucose utilization. ProcNatlAcadSciUSA 91:10625–9.

Perrenoud Q, Geoffroy H, Gauthier B, Rancillac A, Alfonsi F, Kessaris N, Rossier J, Vitalis T, Gallopin T. 2012a. Characterization of Type I and Type II nNOS-Expressing Interneurons in the Barrel Cortex of Mouse. Front Neural Circuits 6:36. doi:10.3389/fncir.2012.00036

Perrenoud Q, Rossier J, Ferezou I, Geoffroy H, Gallopin T, Vitalis T, Rancillac A. 2012b. Activation of cortical 5-HT(3) receptor-expressing interneurons induces NO mediated vasodilatations and NPY mediated vasoconstrictions. Front Neural Circuits 6:50. doi:10.3389/fncir.2012.00050

Price CJ, Cauli B, Kovacs ER, Kulik A, Lambolez B, Shigemoto R, Capogna M. 2005. Neurogliaform neurons form a novel inhibitory network in the hippocampal CA1 area. JNeurosci 25:6775–6786.

Ramos B, Baglietto-Vargas D, del Rio JC, Moreno-Gonzalez I, Santa-Maria C, Jimenez S, Caballero C, Lopez-Tellez JF, Khan ZU, Ruano D, Gutierrez A, Vitorica J. 2006. Early neuropathology of somatostatin/NPY GABAergic cells in the hippocampus of a PS1xAPP transgenic model of Alzheimer’s disease. NeurobiolAging 27:1658–1672.

Rancillac A, Rossier J, Guille M, Tong XK, Geoffroy H, Amatore C, Arbault S, Hamel E, Cauli B. 2006. Glutamatergic Control of Microvascular Tone by Distinct GABA Neurons in the Cerebellum. JNeurosci 26:6997–7006.

Roman RJ. 2002. P-450 metabolites of arachidonic acid in the control of cardiovascular function. Physiol Rev 82:131–185.

Ruff CF, Juarez AF, Dienel SJ, Rakymzhan A, tamirano-Espinoza A, Couey J, Fukuda M, Watson AM, Su A, Fish KN, Rubio ME, Hooks BM, Ross SE, Vazquez AL. 2024. Long-range inhibitory neurons mediate cortical neurovascular coupling. Cell Rep 43:113970.

Rungta RL, Chaigneau E, Osmanski BF, Charpak S. 2018. Vascular Compartmentalization of Functional Hyperemia from the Synapse to the Pia. Neuron 99:362–375.

Rungta RL, Osmanski BF, Boido D, Tanter M, Charpak S. 2017. Light controls cerebral blood flow in naive animals. NatCommun 8:14191.

Sada N, Lee S, Katsu T, Otsuki T, Inoue T. 2015. Targeting LDH enzymes with a stiripentol analog to treat epilepsy. Science 347:1362–1367. doi:10.1126/science.aaa1299

Schaeffer S, Iadecola C. 2021. Revisiting the neurovascular unit. NatNeurosci 24:1198–1209.

Shabir O, Sharp P, Rebollar MA, Boorman L, Howarth C, Wharton SB, Francis SE, Berwick J. 2020. Enhanced Cerebral Blood Volume under Normobaric Hyperoxia in the J20-hAPP Mouse Model of Alzheimer’s Disease. SciRep 10:7518.

Sharp PS, Ameen-Ali KE, Boorman L, Harris S, Wharton S, Howarth C, Shabir O, Redgrave P, Berwick J. 2020. Neurovascular coupling preserved in a chronic mouse model of Alzheimer’s disease: Methodology is critical. J Cereb Blood Flow Metab Off J Int Soc Cereb Blood Flow Metab 40:2289–2303. doi:10.1177/0271678X19890830

Silberberg G, Markram H. 2007. Disynaptic inhibition between neocortical pyramidal cells mediated by Martinotti cells. Neuron 53:735–746.

Slais K, Vorisek I, Zoremba N, Homola A, Dmytrenko L, Sykova E. 2008. Brain metabolism and diffusion in the rat cerebral cortex during pilocarpine-induced status epilepticus. ExpNeurol 209:145–154.

Smith SJ, Sumbul U, Graybuck LT, Collman F, Seshamani S, Gala R, Gliko O, Elabbady L, Miller JA, Bakken TE, Rossier J, Yao Z, Lein E, Zeng H, Tasic B, Hawrylycz M. 2019. Single-cell transcriptomic evidence for dense intracortical neuropeptide networks. Elife 8.

Sun X, Li P, Luo W, Chen S, Feng N, Wang J, Luo Q. 2010. Investigating the effects of dimethylsulfoxide on hemodynamics during cortical spreading depression by combining laser speckle imaging with optical intrinsic signal imaging. Lasers SurgMed 42:649–655.

Taniguchi H, He M, Wu P, Kim S, Paik R, Sugino K, Kvitsiani D, Fu Y, Lu J, Lin Y, Miyoshi G, Shima Y, Fishell G, Nelson SB, Huang ZJ. 2011. A resource of Cre driver lines for genetic targeting of GABAergic neurons in cerebral cortex. Neuron 71:995–1013. doi:10.1016/j.neuron.2011.07.026

Tasic B, Menon V, Nguyen TN, Kim TK, Jarsky T, Yao Z, Levi B, Gray LT, Sorensen SA, Dolbeare T, Bertagnolli D, Goldy J, Shapovalova N, Parry S, Lee C, Smith K, Bernard A, Madisen L, Sunkin SM, Hawrylycz M, Koch C, Zeng H. 2016. Adult mouse cortical cell taxonomy revealed by single cell transcriptomics. NatNeurosci 19:335–346.

Thevenaz P, Ruttimann UE, Unser M. 1998. A pyramid approach to subpixel registration based on intensity. IEEE TransImage Process 7:27–41.

Tomioka R, Okamoto K, Furuta T, Fujiyama F, Iwasato T, Yanagawa Y, Obata K, Kaneko T, Tamamaki N. 2005. Demonstration of long-range GABAergic connections distributed throughout the mouse neocortex. EurJNeurosci 21:1587–1600.

Tricoire L, Kubota Y, Cauli B. 2013. Cortical NO interneurons: from embryogenesis to functions. Front Neural Circuits 7:105.

Tricoire L, Pelkey KA, Daw MI, Sousa VH, Miyoshi G, Jeffries B, Cauli B, Fishell G, McBain CJ. 2010. Common origins of hippocampal ivy and nitric oxide synthase expressing neurogliaform cells. JNeurosci 30:2165–2176.

Turner KL, Gheres KW, Proctor EA, Drew PJ. 2020. Neurovascular coupling and bilateral connectivity during NREM and REM sleep. eLife 9:e62071. doi:10.7554/eLife.62071

Uhlirova H, Kilic K, Tian P, Thunemann M, Desjardins M, Saisan PA, Sakadzic S, Ness TV, Mateo C, Cheng Q, Weldy KL, Razoux F, Vanderberghe M, Cremonesi JA, Ferri CG, Nizar K, Sridhar VB, Steed TC, Abashin M, Fainman Y, Masliah E, Djurovic S, Andreassen O, Silva GA, Boas DA, Kleinfeld D, Buxton RB, Einevoll GT, Dale AM, Devor A. 2016. Cell type specificity of neurovascular coupling in cerebral cortex. Elife 5:e14315.

Vanlandewijck M, He L, Mae MA, Andrae J, Ando K, Del GF, Nahar K, Lebouvier T, Lavina B, Gouveia L, Sun Y, Raschperger E, Rasanen M, Zarb Y, Mochizuki N, Keller A, Lendahl U, Betsholtz C. 2018. A molecular atlas of cell types and zonation in the brain vasculature. Nature 554:475–480.

Viollet C, Simon A, Tolle V, Labarthe A, Grouselle D, Loe-Mie Y, Simonneau M, Martel G, Epelbaum J. 2017. Somatostatin-IRES-Cre Mice: Between Knockout and Wild-Type? Front EndocrinolLausanne 8:131.

Wood J, Garthwaite J. 1994. Models of the diffusional spread of nitric oxide: implications for neural nitric oxide signalling and its pharmacological properties. Neuropharmacology 33:1235–1244.

Xie L, Kang H, Xu Q, Chen MJ, Liao Y, Thiyagarajan M, O’Donnell J, Christensen DJ, Nicholson C, Iliff JJ, Takano T, Deane R, Nedergaard M. 2013. Sleep drives metabolite clearance from the adult brain. Science 342:373–377.

Zeisel A, Manchado AB, Codeluppi S, Lonnerberg P, La MG, Jureus A, Marques S, Munguba H, He L, Betsholtz C, Rolny C, Castelo-Branco G, Hjerling-Leffler J, Linnarsson S. 2015. Cell types in the mouse cortex and hippocampus revealed by single-cell RNA-seq. Science 347:1138–1142.

Zemankovics R, Kali S, Paulsen O, Freund TF, Hajos N. 2010. Differences in subthreshold resonance of hippocampal pyramidal cells and interneurons: the role of h-current and passive membrane characteristics. JPhysiol 588:2109–2132.

Zhang D, Raichle ME. 2010. Disease and the brain’s dark energy. NatRevNeurol 6:15–28.

Zhang D, Ruan J, Peng S, Li J, Hu X, Zhang Y, Zhang T, Ge Y, Zhu Z, Xiao X, Zhu Y, Li X, Li T, Zhou L, Gao Q, Zheng G, Zhao B, Li X, Zhu Y, Wu J, Li W, Zhao J, Ge WP, Xu T, Jia JM. 2024. Synaptic-like transmission between neural axons and arteriolar smooth muscle cells drives cerebral neurovascular coupling. NatNeurosci.

Zielinski MR, Atochin DN, McNally JM, McKenna JT, Huang PL, Strecker RE, Gerashchenko D. 2019. Somatostatin+/nNOS+ neurons are involved in delta electroencephalogram activity and cortical-dependent recognition memory. Sleep 42.

Zonta M, Angulo MC, Gobbo S, Rosengarten B, Hossmann KA, Pozzan T, Carmignoto G. 2003. Neuron-to-astrocyte signaling is central to the dynamic control of brain microcirculation. NatNeurosci 6:43–50.

Zott B, Simon MM, Hong W, Unger F, Chen-Engerer HJ, Frosch MP, Sakmann B, Walsh DM, Konnerth A. 2019. A vicious cycle of beta amyloid-dependent neuronal hyperactivation. Science 365:559–565.

