## Supplementary material for "Bidirectional control of neurovascular coupling by cortical somatostatin interneurons": Supplemntal Table 1 and Figures 1-6

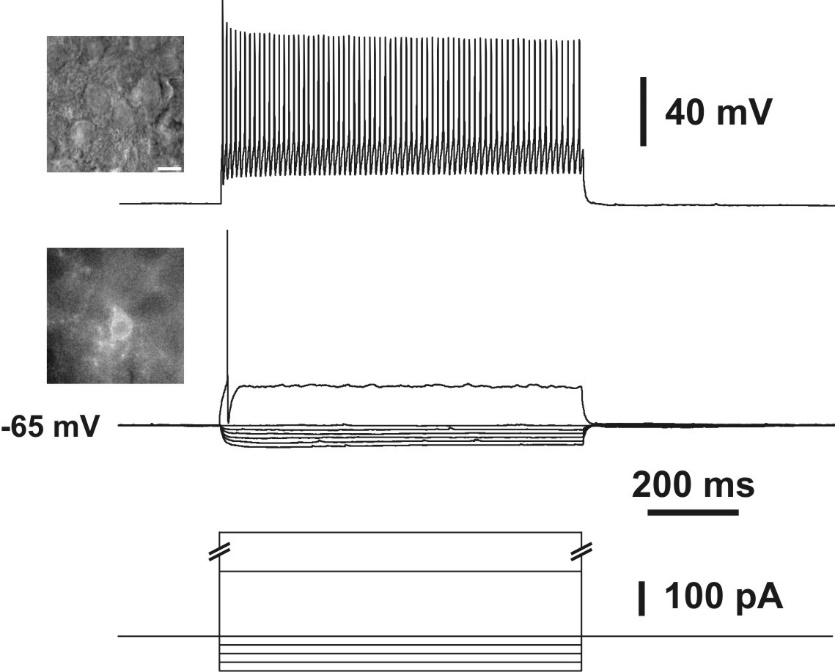

**Supplementary Figure 1: Electrophysiological characterization of a fast spiking Sst-ChR2 neurons.** Recording showing the voltage responses induced by 800 ms current injections (bottom traces) of -100 pA, -80 pA, -60 pA, -40 pA, -20 pA, 0 pA, +190 and +600 pA in a cortical EYFP/ChR2-expressing neuron of a Sst-Cre:Ai32 mouse. Note the small voltage response to hyperpolarizing current injections (middle traces). Just-above threshold current pulse (+190 pA) triggered a discharge of an action potential (middle traces). Note the sharp and fast AHP. Near saturation, a strong depolarizing current (+600 pA) triggered a discharge of action potentials with a weak frequency adaptation and a monotonous amplitude accommodation. Top inset: IR videomicroscopy picture of the recorded neuron, the pial surface is upward (scale bar, 20 µm). Bottom insets, corresponding field of view showing the EYFP fluorescence of the recorded neuron.

**Supplementary Table 1. Electrophysiological properties of Sst-ChR2 interneuron**

|  | **Adapting-*Sst***  **(n= 15)** | **Fast Spiking**  **(n=1)** |
| --- | --- | --- |
| **Passive properties** | | |
| **(1) Resting potential (mV)** | -59.5 ± 4.5 | -58.0 |
| **(2) Input resistance (MΩ)** | 608 ± 231 | 129 |
| **(3) Time constant (ms)** | 39.1 ± 15.3 | 11.7 |
| **(4) Membrane capacitance (pF)** | 68.5 ± 31.1 | 90.7 |
| **(5) Sag index (%)** | 21.2 ± 10.6 | 10.5 |
| **Just above threshold properties** | | |
| **(6) Rheobase (pA)** | 8 ± 14 | 187 |
| **(7) First spike latency (ms)** | 130.1 ± 162.8 | 17.0 |
| **(8) Adaptation (Hz/s)** | -2.6 ± 6.9 | -38.5 |
| **(9) Minimal frequency (Hz)** | 9.3 ± 6.7 | 41.4 |
| **Firing properties** | | |
| **(10) Accommodation (mV)** | 0.7 ± 0.7 | 0.2 |
| **(11) Amplitude of early adaptation (Hz)** | 27.1 ± 10.0 | 30.4 |
| **(12) Time constant of early adaptation (ms)** | 60.5 ± 23.8 | 15.2 |
| **(13) Late adaptation (Hz/s)** | -10.3 ± 9.4 | -21.7 |
| **(14) Maximal frequency (Hz)** | 46.7 ± 16.6 | 98.6 |
| **Action potentials properties** | | |
| **(15) 1^st^ spike amplitude (mV)** | 83.6 ± 8.8 | 85.9 |
| **(16) 1^st^ spike duration (ms)** | 1.3 ± 0.3 | 0.6 |
| **(17) 2^nd^ spike amplitude (mV)** | 79.2 ± 9.1 | 78.3 |
| **(18) 2^nd^ spike duration (ms)** | 1.4 ± 0.3 | 0.7 |
| **(19) Amplitude Reduction (%)** | 5.4 ± 3.9 | 8.9 |
| **(20) Duration Increase (%)** | 3.3 ± 3.8 | 6.3 |
| **AHP and ADP properties** | | |
| **(21) 1^st^ spike fast AHP (mV)** | -18.6 ± 5.5 | -26.5 |
| **(22) 1^st^ spike ADP (mV)** | 2.2 ± 2.7 | 0.0 |
| **(23) 1^st^ spike medium AHP (mV)** | -7.5 ± 6.5 | 0.0 |
| **(24) 1^st^ spike fast AHP latency (ms)** | 5.0 ± 1.4 | 2.6 |
| **(25) 1^st^ spike ADP latency (ms)** | 8.5 ± 7.9 | 0.0 |
| **(26) 1^st^ spike, medium AHP latency (ms)** | 18.5 ± 18.2 | 0.0 |
| **(27) 2^nd^ spike fast AHP (mV)** | -19.4 ± 6.0 | -28.6 |
| **(28) 2^nd^ spike ADP (mV)** | 2.0 ± 2.6 | 0.0 |
| **(29) 2^nd^ spike medium AHP (mV)** | -8.2 ± 7.2 | 0.0 |
| **(30) 2^nd^ spike, fast AHP latency (ms)** | 5.2 ± 1.4 | 2.8 |
| **(31) 2^nd^ spike ADP latency (ms)** | 8.7 ± 9.0 | 0.0 |
| **(32) 2^nd^ spike, medium AHP latency (ms)** | 19.5 ± 19.0 | 0.0 |

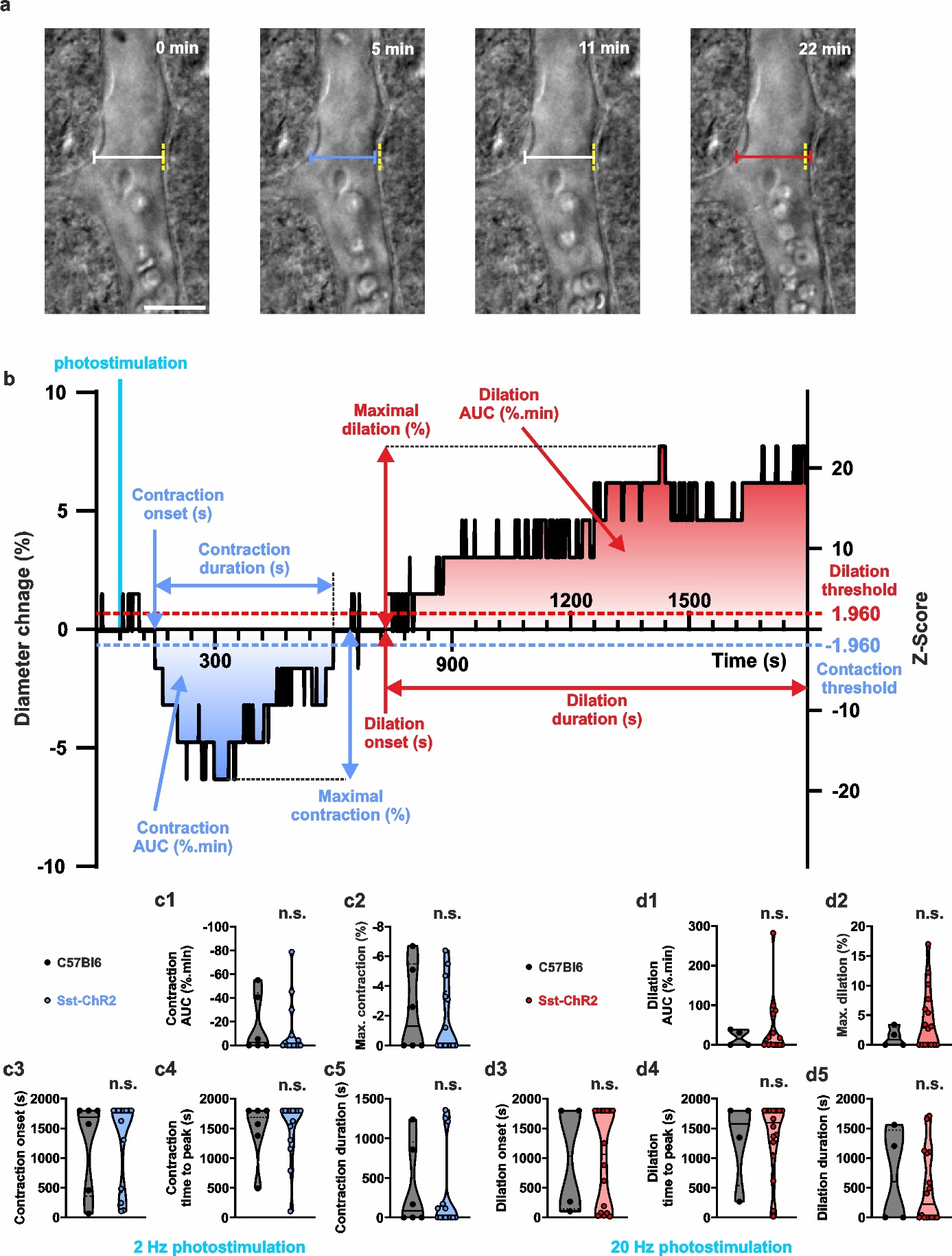

**Supplementary Figure 2: Measurement of vascular response parameters. (a)** Infrared pictures of a diving arteriole before (0 min) and after 20 Hz photostimulation showing a vasoconstriction (5 min) followed by a vasodilation (22 min). The pial surface is upward. Scale bars 20 µm. White, red and blue calipers correspond to the baseline, dilated, and contracted arteriolar diameter, respectively. The dashed yellow lines indicate the initial diameter. **(b)** Kinetics of the vascular response of the arteriole shown in **(a)** expressed as diameter changes (left axis) and Z-score (right axis) relative to the baseline period. The dashed horizontal blue and red lines depict the statistical (p<0.05) Z-score thresholds for contraction and dilation, respectively. The measured parameters include the onset, maximum amplitude, duration, and area under the curve of vascular responses. **(c1-5, d1-5)** Violin plots summarizing the comparison of magnitude of contraction **(c1)** and dilation**(d1)**, maximal contraction **(c2)** and dilation **(d2)**, onset **(c3,d3),** time to peak **(c4,f4)**, and duration **(c5,f5)** of contraction **(c1-5)** and dilation **(d1-5)** between arterioles from slices of Sst-Cre:Ai32 and C57Bl6 mice stimulated at 2 (**c1-5**) and 20 Hz (**d1-5**), respectively. Note that the vasoconstriction and vasodilation, respectively elicited by 2 and 20 Hz stimulation of Sst-ChR2 slices, do not differ from the sporadic vascular changes observed in C57Bl6 slices. Data are presented as individual values. Solid and dashed black bars correspond to median and quartile values. n.s.: not statistically significant.

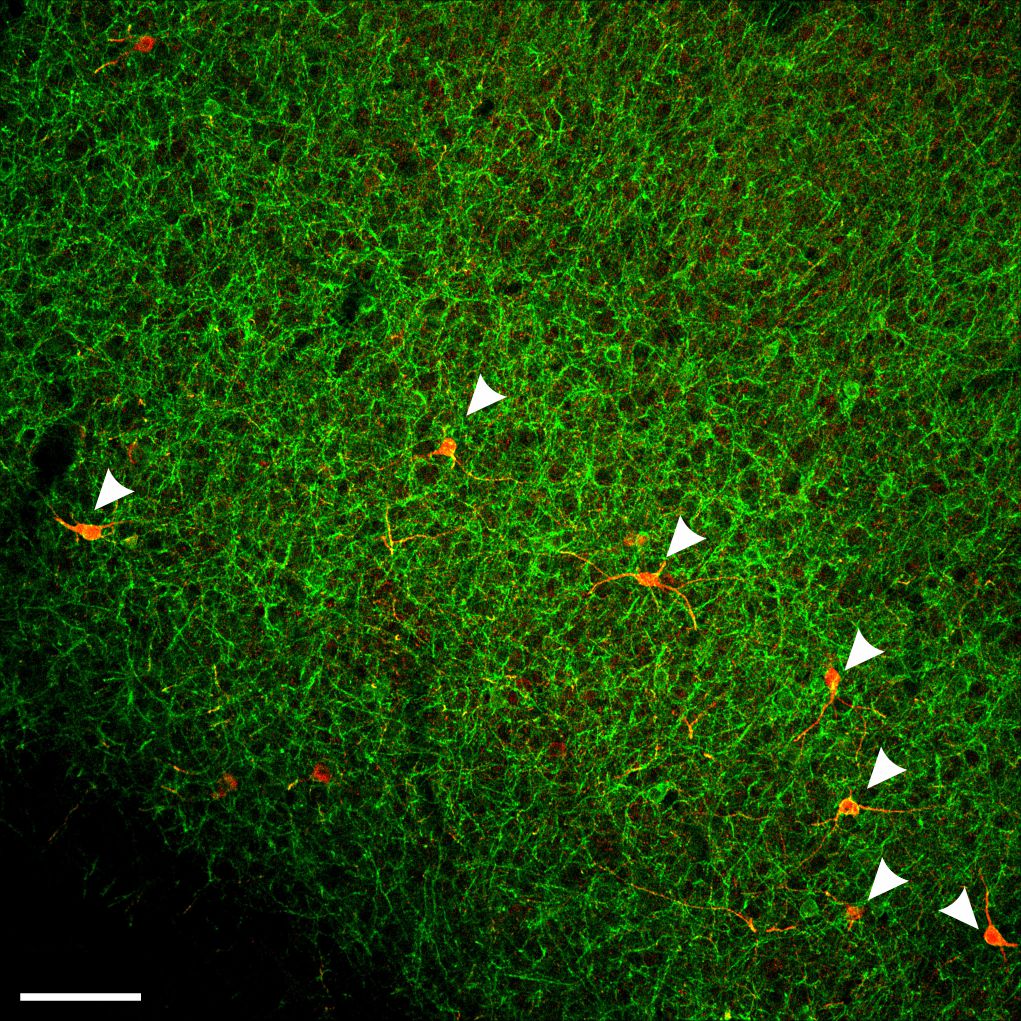

**Supplementary Figure 3: Expression of NOS-1 in Sst-ChR2 neurons**. Single-plane confocal image of double-fluorescence staining showing the expression of ChR2-EYFP (green) and NOS-1 (red) in a cortical slice from a Sst-ChR2 mouse. Note the densely labeled ChR2-EYFP positive fibers and perikarions. Arrowheads indicate ChR2-EYFP-positive neurons immunolabelled for NOS-1. Scale bar: 100 µm.

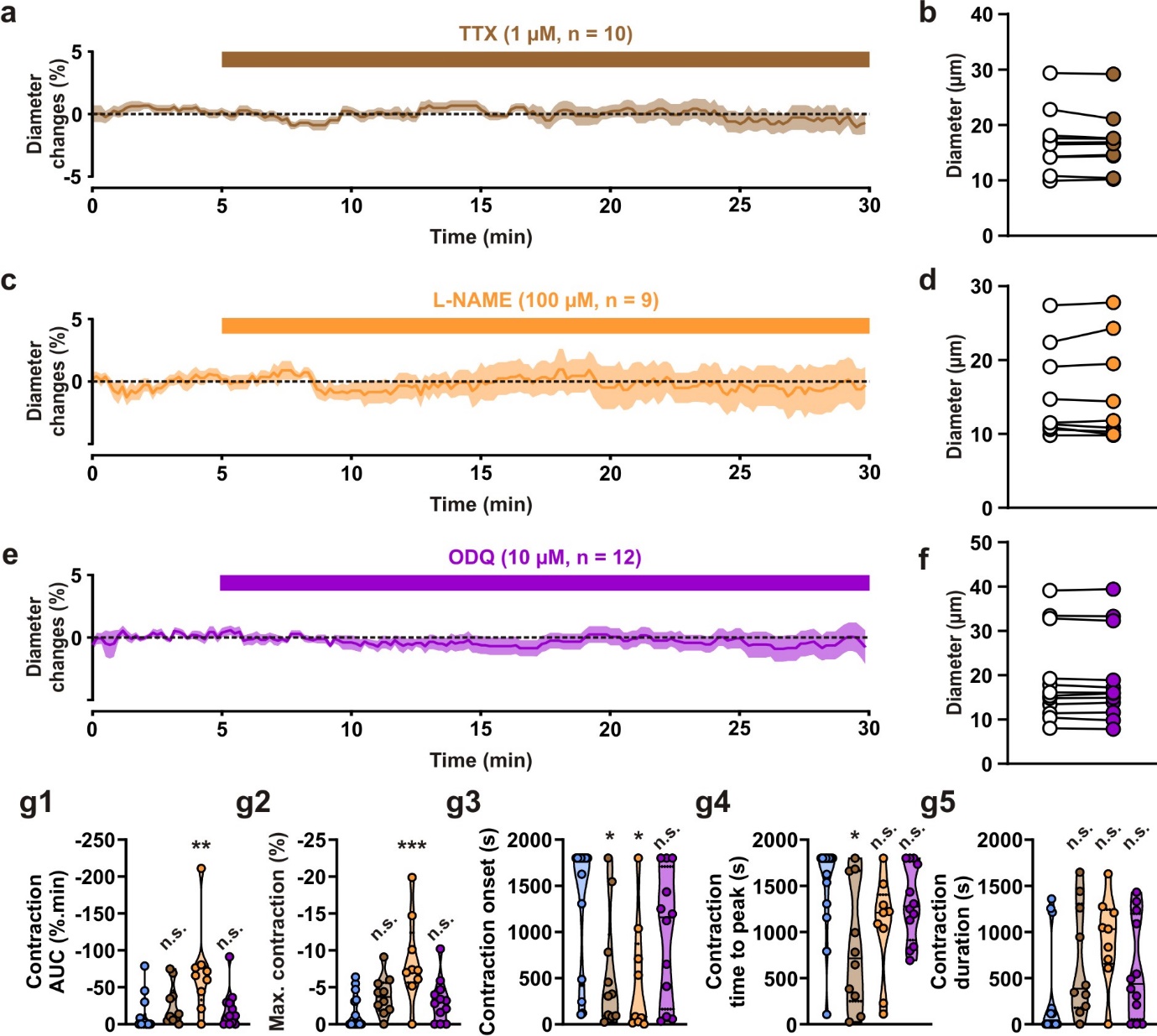

**Supplementary Figure 4: Effects of pharmacological treatments on the basal vascular tone and the vasoconstriction evoked by 2 Hz stimulation.** **(a, c, e)** Kinetics of vascular diameter changes during pharmacological application. Note the absence of diameter changes during the application of 1 µM TTX (**a**, brown horizontal zone, n= 10 arterioles from 6 mice), 100 µM L-NAME (**c**, blue horizontal zone, n= 9 arterioles from 6 mice) and 10 µM ODQ (**e**, purple horizontal zone, n= 12 arterioles from 7 mice). **(b, d, f)** Comparisons of arteriolar diameters during the 5-minute baseline period and the last 5 minutes of the 30-minute pharmacologic treatments. Note the absence of statistically significant differences after TTX **(b)**, L-NAME **(d)** and ODQ **(f)** applications. Violin plots summarizing the effects of TTX (brown), L-NAME (orange) and ODQ (purple) on the magnitude **(g1**, H_(5,51)_= 13.053, p= 0.01102**),** maximum **(g2**, H_(5,51)_= 14.761, p= 0.00522**)**, onset **(g3**, H_(5,51)_= 11.795, p= 0.01894**),** time to peak **(g4**, H_(5,51)_= 10.957, p= 0.02705**)**, and duration **(g5**, H_(5,51)_= 9.278, p= 0.05452**)** of vasoconstriction induced by 2 Hz photostimulation. Note the stronger vasoconstriction under L-NAME treatment. Data are presented as individual values. Solid and dashed black bars correspond to median and quartile values. * , ** and *** statistically different from control condition 2 Hz Sst-ChR2 (blue) with p<0.05, 0.01 and 0.001, respectively.

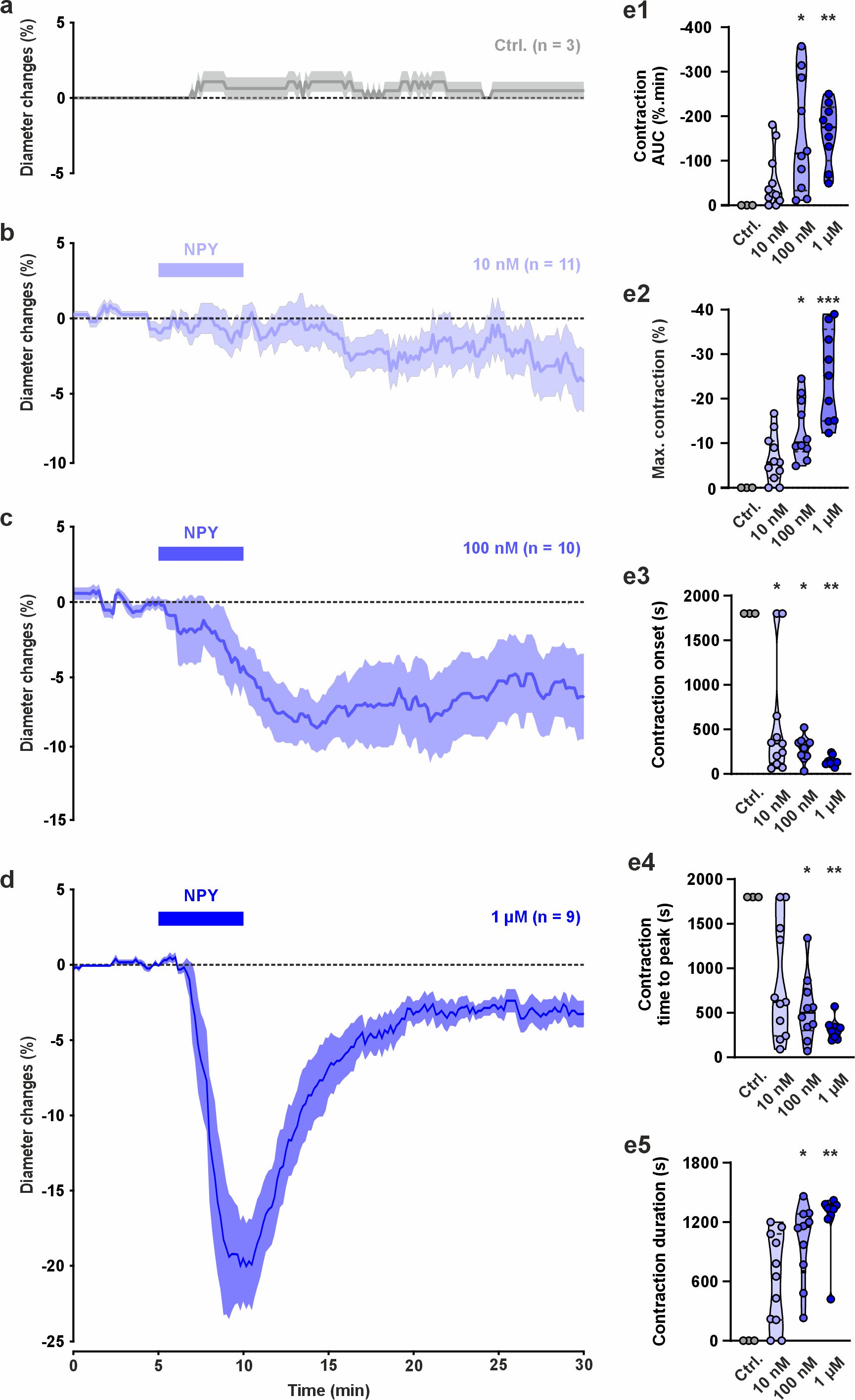

**Supplementary Figure 5: The dose-response effects of NPY-induced vasoconstriction. (a-d)** Kinetics of diameter changes in absence (**a**, gray, n= 3 arterioles from 3 mice) and in presence of NPY applied for 5 minutes at 10 nM (**b**, light blue, n= 11 arterioles from 7 mice), 100 nM (**c**, medium blue, n= 10 arterioles from 8 mice) and 1 µM (**d**, dark blue, n= 9 arterioles from 4 mice). **(b-d)**. **(e1-3, f1-3).** The SEMs envelope the mean traces. **(e1-5)**. Violin plots summarizing the effects of 10 nM (light blue), 100 nM (medium blue) and 1 µM NPY (dark blue) on the on the magnitude **(e1**, H_(4,33)_= 14.474, p= 0.00233**),** maximum **(e2**, H_(4,33)_= 20.060, p= 0.00016**)**, onset **(e3**, H_(4,33)_= 11.740, p= 0.00833**),** time to peak **(e4**, H_(4,33)_= 11.371, p= 0.00988**)**, and duration **(e5**, H_(4,33)_= 17.733, p= 0.00050**)** of vasoconstriction compared to the control condition. Note the lack of consistent diameter changes in the absence of NPY **(a,e)** and the higher magnitude **(e1)** and amplitude **(e2)**, the earlier onset **(e3)** and time to peak **(e4)**, and the longer **(e5)** vasoconstriction with increasing NPY concentrations **(b-d)**. Data are presented as individual values. Solid and dashed black bars correspond to median and quartile values. *, ** and *** statistically different from the control condition with p < 0.05, 0.01 and 0.001, respectively.

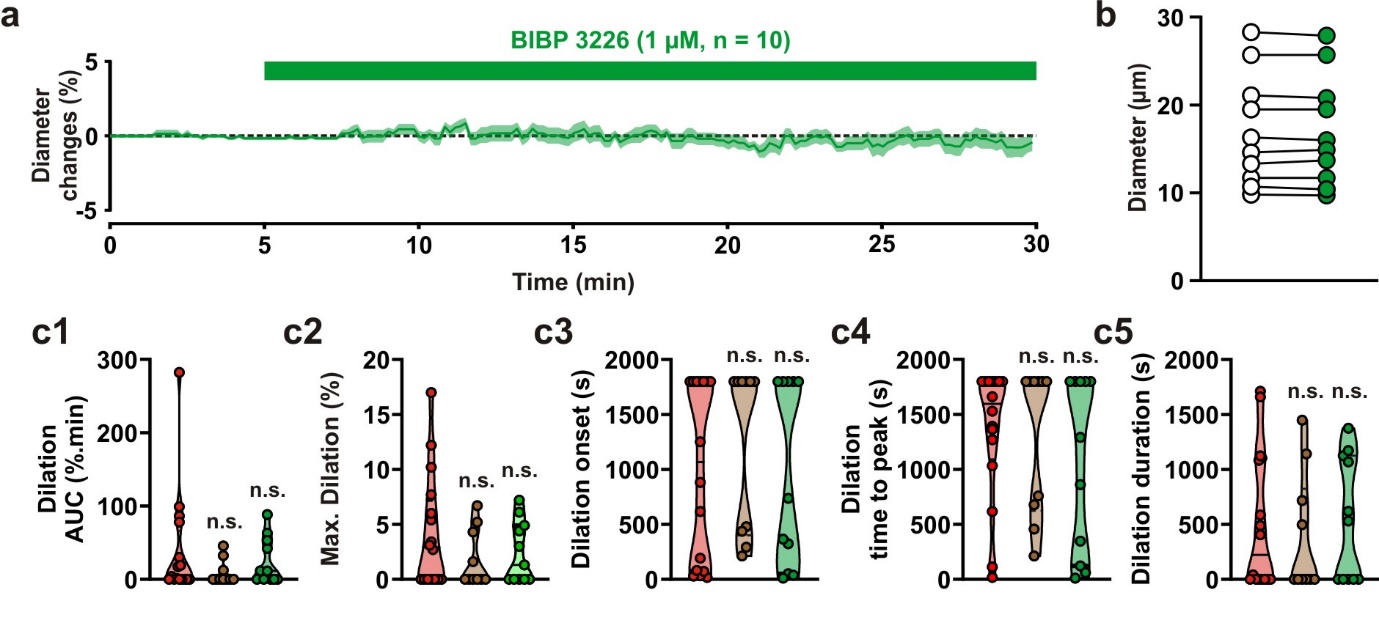

**Supplementary Figure 6: Effects of pharmacological treatments on the basal vascular tone and the vasodilation evoked by 20 Hz stimulation.** **(a)** Kinetics of vascular diameter changes during BIBP3226 application. Note the absence of diameter changes during the application of 1 µM BIBP 3226 (**a**, green horizontal zone, n= 11 arterioles from 9 mice). **(b)** Comparisons of arteriolar diameters during the 5-minute baseline period and the last 5 minutes of the 30-minute BIBP3226 treatment. Note the absence of statistically significant differences after BIBP 3226. applications. Violin plots summarizing the effects of TTX (brown) and BIBP3226 (green) on the magnitude **(c1***,* H_(4,41)_= 1.947, p= 0.58347**),** maximum **(c2***,* H_(4,41)_= 1.714 , p= 0.63375**)**, onset **(c3***,* H_(4,41)_= 1.401 , p=0.70520 **),** time to peak **(c4***,* H_(4,41)_= 0.815, p= 0.84585**)**, and duration **(c5***,* H_(4,41)_= 0.747 , p= 0.86209**)** of vasodilation induced by 20 Hz photostimulation. Neither TTX or BIBP 3226 alter the vasodilation. Data are presented as individual values. Solid and dashed black bars correspond to median and quartile values. n.s. not statistically different from the control condition 20 Hz Sst-ChR2 control condition (red).
